# High density genomic surveillance and risk profiling of clinical *Listeria monocytogenes* subtypes in Germany, 2018-2021

**DOI:** 10.1101/2024.02.16.580625

**Authors:** Sven Halbedel, Sabrina Wamp, Raskit Lachmann, Alexandra Holzer, Ariane Pietzka, Werner Ruppitsch, Hendrik Wilking, Antje Flieger

**Affiliations:** FG11 Division of Enteropathogenic bacteria and Legionella, Consultant Laboratory for Listeria, Robert Koch Institute, Burgstrasse 37, 38855 Wernigerode, Germany; Institute for Medical Microbiology and Hospital Hygiene, Otto von Guericke University Magdeburg, Leipziger Strasse 44, 39120 Magdeburg, Germany; FG35 – Division for Gastrointestinal Infections, Zoonoses and Tropical Infections, Robert Koch Institute, Seestrasse 10, 13353, Berlin, Germany; Austrian Agency for Health and Food Safety, Institute for Medical Microbiology and Hygiene, Beethovenstraße 6, A-8010 Graz, Austria; Austrian Agency for Health and Food Safety, Institute for Medical Microbiology and Hygiene, Währingerstrasse 25a, A-1090 Vienna, Austria

**Author notes:** Corresponding authors, Robert Koch Institute, FG11 Division of Enteropathogenic bacteria and *Legionella*, Burgstrasse 37, D-38855 Wernigerode, Germany, Robert Koch Institute, FG11 Division of Enteropathogenic bacteria and *Legionella*, Burgstrasse 37, D-38855 Wernigerode, Germany. Shared first authors (Sven Halbedel, Sabrina Wamp).

**Keywords:** epidemiology, outbreak, *inlF*, *flaR*, *clpP1*

## Abstract

Foodborne infections represent a significant public health concern, particularly when outbreaks affect many individuals over prolonged time. Systematic collection of pathogen isolates from infected patients, whole genome sequencing and phylogenetic analyses allow recognition and termination of outbreaks after source identification and risk profiling of abundant lineages. We here present a multi-dimensional analysis of >1,800 genome sequences from clinical *L. monocytogenes* isolates collected in Germany between 2018-2021. These isolates covered 62% of all notified cases and belonged to 188 infection clusters. 42% of these clusters were active for >12 months, 60% generated cases cross-regionally, including 11 multinational clusters. 37% of the clusters were caused by sequence type (ST) ST6, ST8 and ST1 clones and for selected clusters, we provide further epidemiological and genetic information. Frequencies of materno-fetal and brain infections allowed risk profiling of the most abundant STs, differentiating ST1 as hyper- and ST8, ST14, ST29 as well as ST155 as hypovirulent from ST6 with average virulence potential. Hepatocyte infection experiments confirmed these virulence differences. Inactivating mutations were found in several virulence and house-keeping genes, particularly in hypovirulent STs. Our work supports prioritization of clusters for epidemiological investigations and reinforces the need to analyse the mechanisms underlying hyper- and hypovirulence.

## Introduction

Listeriosis is a severe foodborne infection and may arise when food contaminated by the Gram-positive bacterium *Listeria monocytogenes* is consumed. Even though contamination of food items and consequently pathogen exposure is quite common (1), listeriosis generally occurs with low frequency, as reflected by the annual incidence of notified cases, which is in the range of 0.2-0.9 patients per 100,000 inhabitants in Europe and North America (2–4). The immune system ensures efficient pathogen clearance after crossing of the gut epithelium (5), explaining low incidence of symptomatic cases in otherwise healthy individuals, where the infection may occur as asymptomatic or self-limiting gastroenteritis. However, invasive disease with manifestations such as septicemia, neurolisteriosis or maternal-fetal infections with high lethality may develop in immunocompromised patients or pregnant women and neonates, respectively (6). Case fatality rates between 13-46% have been reported, depending on the length of the patient observation interval after infection or on the various disease manifestations (2, 7, 8). Such rates are exceptionally high and generally not observed with other bacterial gastrointestinal pathogens (7, 9).

Prevention of *L. monocytogenes* food contamination is challenging, since the bacterium is ubiquitously found in many environmental habitats as well as in the digestive tract and lymphatic organs of productive livestock, leading to frequent contamination of raw materials or to cross-contamination in food processing plants (10–12). Contamination control strategies are implemented at different levels of the food production chain and include optimization of farming practices, improvement of product sanitation and storage conditions or implementation of cleaning and disinfection protocols in the production environment (13–15). Moreover, European legislation has defined contamination limits for *L. monocytogenes* in ready-to-eat or infant food (16). While all these measures doubtlessly help to keep listeriosis incidence low, large and protracted outbreaks of listeriosis regularly occur (17–19), highlighting the importance of disease surveillance systems for detection of clusters of epidemiologically linked cases over a prolonged time as a prerequisite to identify and inactivate the underlying source.

During the last years, several countries or supranational entities such as the European Union have established pathogen surveillance systems based on whole genome sequencing (WGS) of clinical and food isolates to allow cluster detection and assignment of food sources (20–24). As a result, listeriosis outbreaks can now be detected in real time, enabling food authorities to implement countermeasures during ongoing outbreaks. WGS has also generated new insights in the relative contribution of different food vectors to the disease burden. We have shown this in a recent WGS based surveillance study, where we estimated that ∼30% of all German listeriosis cases with a known food source are related to consumption of salmon products (25). Moreover, our knowledge on genomic diversity of *L. monocytogenes* strains, including the identification of hypo- and hypervirulent subtypes (26), their genetic determinants and the description of novel markers associated with stress, biocide and antibiotic resistance has strongly benefited from systematic genome sequencing (27–29).

In Germany, incidence of listeriosis steadily increased from 0.4/100,000 in 2011 to 0.9/100,000 in 2017, but was slightly lower in subsequent years (2). This recent trend of declining case numbers coincided with the introduction of WGS in the German listeriosis surveillance system in 2018 (20), which had led to the detection and termination of large listeriosis outbreaks (18, 30–32). We here present a high-density analysis of the genomic diversity and population structure of clinical *L. monocytogenes* strains isolated from listeriosis patients in Germany between 2018-2021 including a risk profiling of subtypes with international importance.

## Results

### Bipartite population structure of clinical *L. monocytogenes* isolates from Germany

In Germany, the detection of *L. monocytogenes* in primary sterile clinical specimens and in swabs from newborns has to be notified to public health authorities, while the bacterial isolates can be sent to the Consultant Laboratory (CL) for *Listeria* at the Robert Koch Institute for further analysis on a voluntary basis. Between 2018-2021, 2,464 cases of human listeriosis were notified in Germany and 1,802 clinical *L. monocyctogenes* isolates were received by the CL. 1,538 of the 1,802 isolates (85%) were allocated to a notified case, while 264 isolates (15%) were sent to the CL but specific allocation to a notified case was not possible. Thus, *L. monocytogenes* isolates were available and assignable for 62% of the notified listeriosis cases in Germany. The portion of notification cases with accompanying isolate submissions ranged from 26-74% depending on the federal country (Fig. 1).

**Figure 1:**
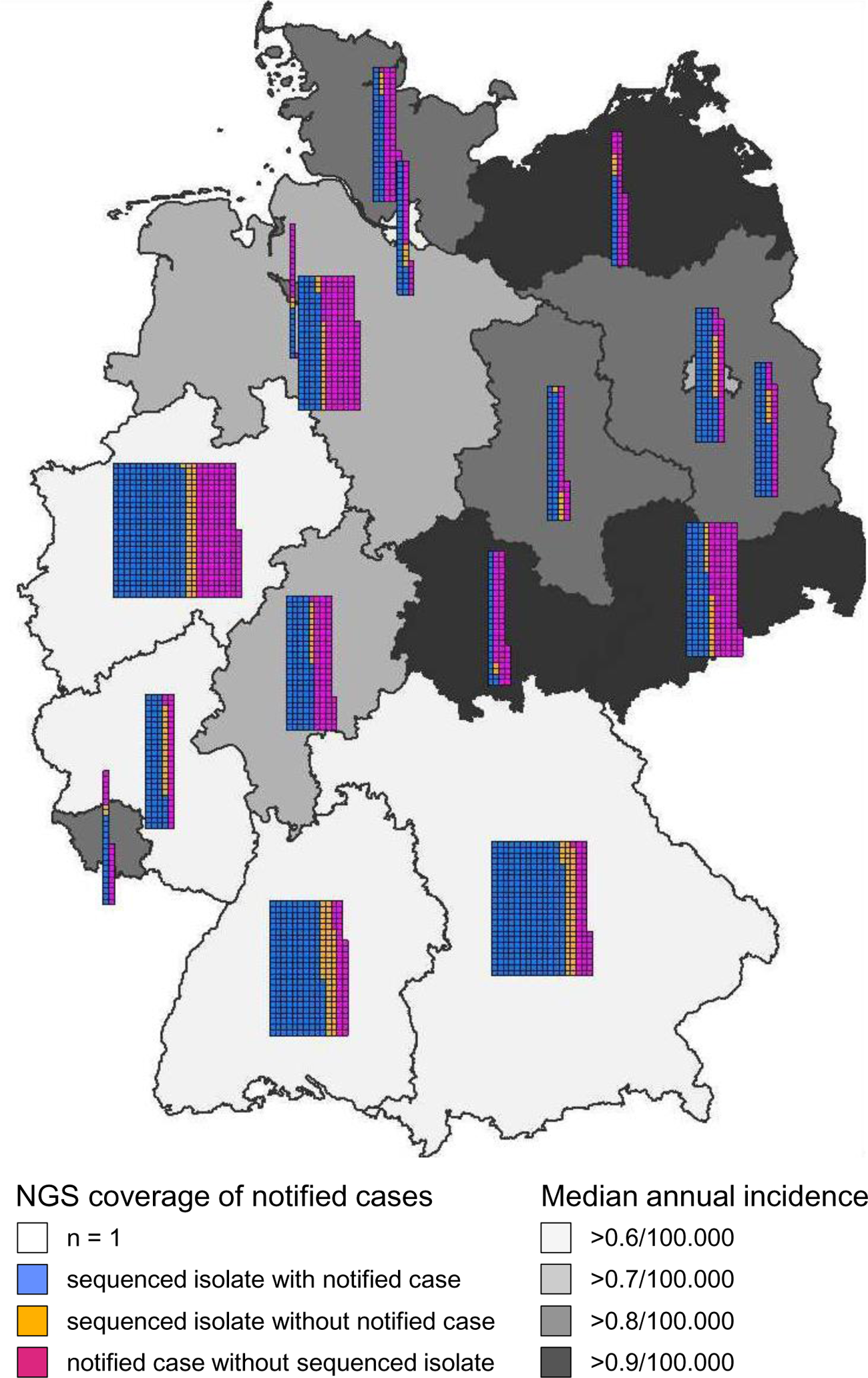
WGS coverage of human listeriosis cases in Germany, 2018-2021. Geographic origin of 2,464 notified listeriosis cases and 1,802 isolate submissions in Germany during 2018-2021. The numbers of notified cases that were accompanied by isolate submissions, the numbers of notified cases without isolate submissions and of isolate submissions for which no notification was initiated are shown for each of the 16 German federal countries. The median incidence per year for the 2018-2021 period is also shown for each federal country.

The genomes of all *L. monocytogenes* strains were sequenced and molecular PCR serogroups were extracted from sequencing data. The majority of the isolates belonged to molecular PCR serogroups IVb (n=830; 46%), IIa (n=781; 43%) and IIb (n=164; 9%), whereas only few isolates were of serogroup IIc (n=14), IVbv-1 (n=10), IVa (n=2) and IVc (n=1) (Fig. 2). Thus, the population primarily is made up of isolates belonging to phylogenetic lineages I and II (Fig. 2), while isolates belonging to other lineages are not found or were underrepresented. Multi locus sequence typing (MLST) showed that sequence types (ST) ST6 (n=300; 17%), ST1 (n=266; 15%), ST8 (n=209; 12%), ST2 (n=91; 5%) and ST451 (n=76; 4%) were the five most prevalent STs (Fig. 3A). The high prevalence of ST6, ST1, ST8 among clinical isolates in Germany had already been observed in a previous study on German isolates from 2007-2017 (20) and isolates of these subtypes are found globally just as the other most prevalent German STs shown in Fig. 3A. In total, the 1,802 isolates grouped into 109 different MLST STs in 59 MLST clonal complexes (CCs) (Fig. 2, Tab. S1). For 5 STs an CC number has not been defined yet.

**Figure 2:**
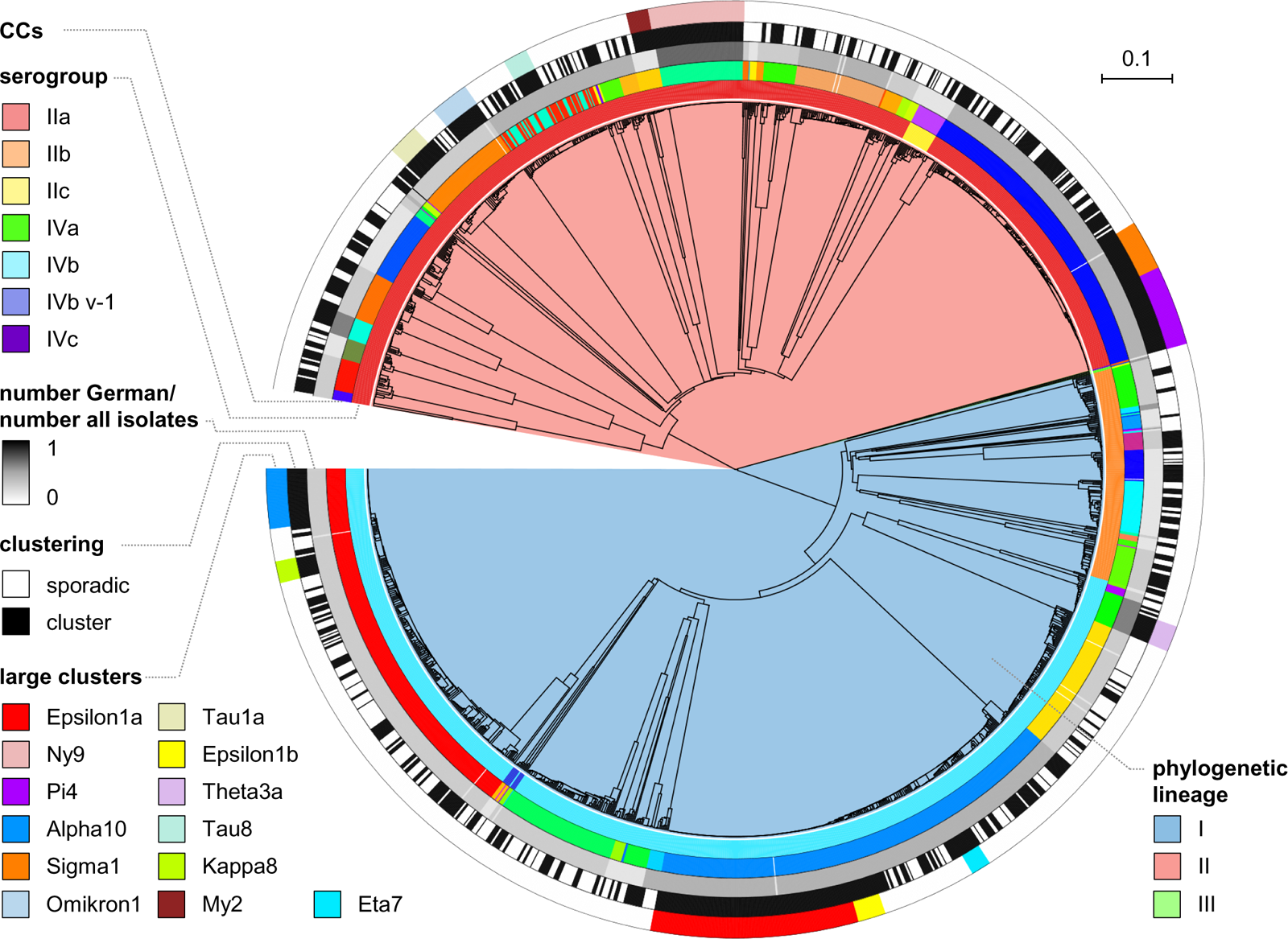
WGS-based subtyping of clinical *L. monocytogenes* strains from Germany. UGPMA tree showing population structure of 1,802 clinical *L. monocytogenes* isolates collected between 2018-2021 in Germany. The tree was calculated from 1,701 locus cgMLST data. Phylogenetic lineages, molecular serogroups, 7 locus MLST STs and attribution to an outbreak cluster are indicated by different color codes. The largest outbreaks in this period are highlighted at the outmost circle. International spread of the STs observed in Germany is expressed as the number of German isolates divided by the number of all (German + known non-German) isolates for each ST (grey shaded ring).

**Figure 3:**
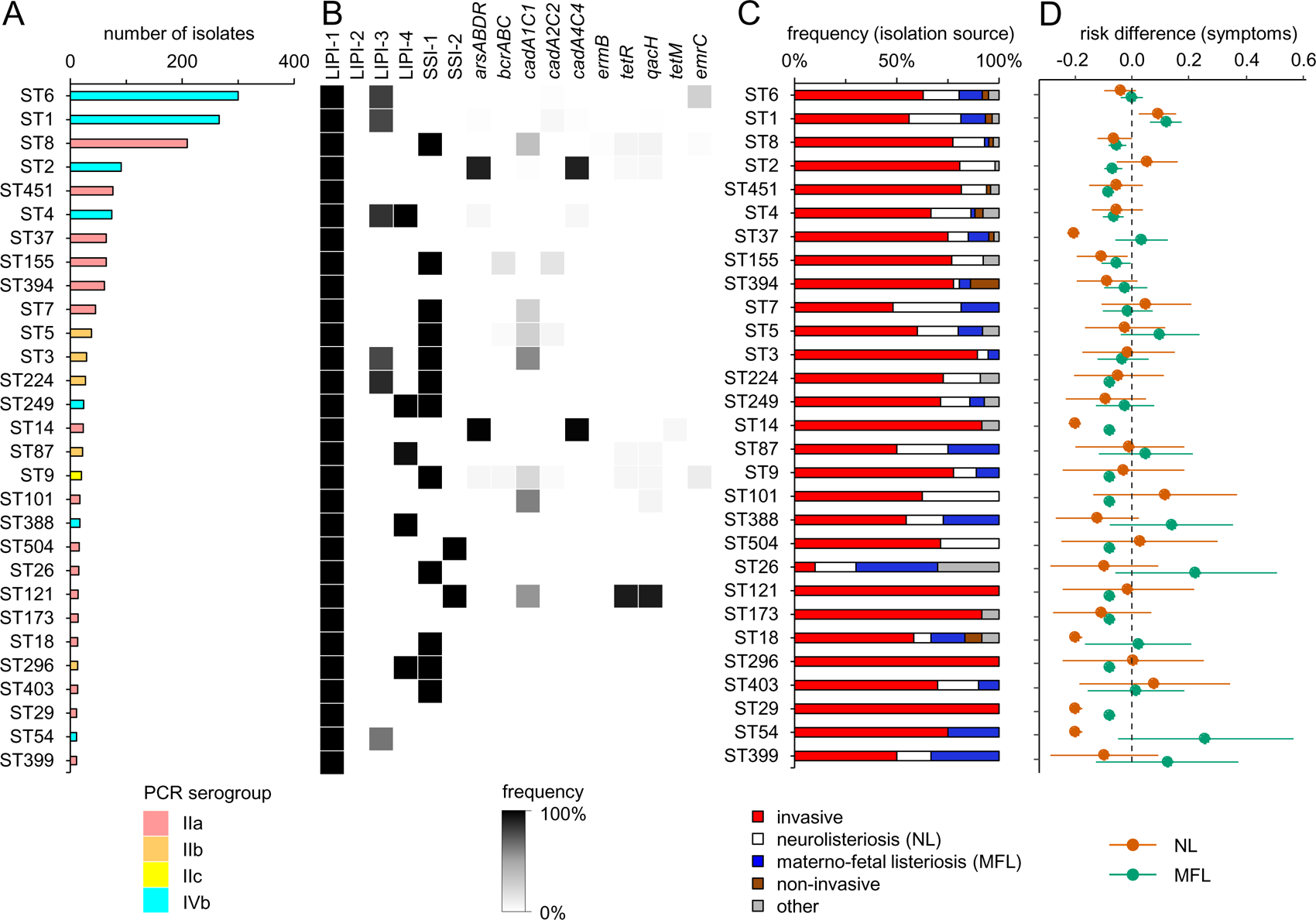
Genomic and clinical characteristics of the most prevalent *L. monocytogenes* STs. (A) Distribution of sequence types among the collection of German *L. monocytogenes* isolates. Only STs with more than 10 isolates are shown. Bars are coloured according to their molecular serogroup. (B) Presence of the four *L. monocytogenes* pathogenicity islands (LIPI), the two stress survival islets (SSI) and selected resistance genes within the different STs. Grey shading reflects the prevalence of a given marker within all analysed isolates of the respective ST. (C) Clinical disease manifestation as reported during isolate submission to the consultant laboratory. For 1,113 out of the 1,802 isolates information on the disease manifestation was available and grouped into five different categories. (D) Clinical disease manifestation as reported during case notification. Risks for NL and MFL manifestations are expressed as risk differences together with 95% confidence intervals. Abbreviations: NL – neurolisteriosis, MFL – materno-fetal listeriosis.

### Comparison with global clinical *L. monocytogenes* isolates identifies autochthonous STs

We wondered whether there are major genomic differences between clinical *L. monocytogenes* in Germany and isolates collected in other countries. In order to address this, we downloaded all genomes of clinical *L. monocytogenes* isolates with non-German origin that had been deposited on the NCBI pathogen detection server until 21^st^ September 2022. This included 15,156 genomes of strains that were isolated from 48 different countries from all world regions (Fig. S1A), however, isolate genomes from the United States (n=7,899) and the United Kingdom (n=1,568) were the most frequent (Fig. S1B). 959 different MLST STs were assigned to the clinical *L. monocytogenes* genomes with non-German origin.

Comparison of the ST assignments for isolates from in-and outside Germany showed that the majority of STs/CCs observed in Germany (80% of the German STs covering 98.5% of the German isolates) were also reported to cause disease outside Germany (Fig. 2). Likewise, German isolates cover most of the phylogenetic branches of pathogenic *L. monocytogenes* from locations outside the country (Fig. S1C), suggesting a cross-country distribution of pathogenic *L. monocytogenes* STs in general. However, an overrepresentation of ST249 isolates (CC315) was observed among the clinical 2018-2021 strains from Germany compared to the international sequences, as 24 clinical ST249 isolates have been collected in Germany and only one is known from outside Germany (Slovenia). Likewise, 14 clinical ST173 isolates from Germany, but only 7 from locations outside Germany, in particular from the Netherlands and the United Kingdom were detected. While the German and the Slovenian ST249 isolates are not closely related, all ST173 isolates belong to the same cgMLST cluster (My2, Tab. S1), reflecting a cross-country outbreak (25). Thus, ST249 and ST173 show limited geographic distribution and therefore may represent autochthonous clones in Germany. In contrast, 868 STs, which are associated with disease outside Germany, are not present in the German 2018-2021 collection. Of these, ST321, ST378 and ST389, predominantly from the United States, New Zealand and/or Taiwan, have the largest number of non-German isolate genomes available at the NCBI server.

### Virulome and resistome differences among frequent clinical subtypes

Allele calling showed that *Listeria* pathogenicity island 1 (LIPI-1) was present in the genomes of all German isolates, while LIPI-2, typically present in *L. ivanovii* (33) or as a truncated version in the novel hybrid sub-lineage HSL-II (34) was not found in any of them (Fig. 3B). For the STs with >10 isolates, LIPI-3, encoding listeriolysin S (35), was detected in ST1, ST3, ST4, ST6 and ST224 isolates as well as in approximately half of the ST54 isolates, corresponding to 35% of all isolates. LIPI-4 associated with neuro-invasion (26) was found in ST4, ST87, ST249, ST296 and ST388 isolates (11% of all isolates). Thus, ST4 was the only group among the more frequent STs that contained three of the four known pathogenicity islands (Fig. 3B), while LIPI-3 and LIPI-4 were generally absent from lineage II and III isolates.

Stress survival islet SSI-1 supporting growth at low pH and high salt conditions (36) was found in 31% of the isolates belonging to different STs, while SSI-2 promoting growth under alkaline and oxidative stress (27) was present only in ST121 and ST504 strains (2% of all isolates). We also observed a tight association of ST2 (92%) and ST14 strains (>98%) with the *arsABDR* and *cadA4/cadC4* resistance genes encoding heavy metal resistance determinants (37, 38), and of ST121 strains with the *qacH* and *tetR* genes (93%) for resistance against quaternary ammonium compounds (39) (Fig. 3B). Taken together, despite their high isolation frequency of ST6, ST1 and ST8 isolates, they are not characterized by a higher number of pathogenicity islands, stress or resistance genes, suggesting that other factors explain their predominance.

### Analysis of clinical metadata highlights ST1 isolates as hypervirulent

Information on the disease manifestation and/or isolation source accompanied isolate submissions for 1,113 of the 1,802 isolates (62%). Specifically, 69% of the isolates were from invasive listeriosis cases (isolation from blood, ascites, synovial, pleural or lymph fluid, abscesses, wounds and histologic specimens or from patients suffering from sepsis, bacteremia or fever), 17% from invasive neurolisteriosis (NL) patients (isolation from cerebrospinal fluid or from patients with meningitis or encephalitis), 7% from maternal-fetal listeriosis (MFL), 2% from non-invasive conditions (stool) and 5% from other manifestations. Remarkably, there were no ST14, ST29, ST121, ST173 and ST296 isolates from MFL or NL samples according to this set of data (Fig. 3C). To further analyse this, we determined the risk differences for MFL or NL associated with infections caused by different *L. monocytogenes* STs using notification data. This data is based upon reported details from the local health authorities on disease manifestation and/or symptoms. Information on the establishment of MFL (n=98 cases), NL (n=203) or not was available for 1,323 isolate/notification case pairs in total. ST6 (serogroup IVb) served as a reference in the risk difference calculations because of its high prevalence. Compared to ST6, risks for MFL and NL were both reduced for infections caused by ST8, ST14, ST29 and ST155 strains (all belonging to serogroup IIa) but increased for infections caused by ST1 strains (serogroup IVb, Fig. 3D). Thus, these STs were here referred to as hypo-and hypervirulent, respectively. We further noticed that several STs were associated with reduced risks to establish MFL (ST2, ST4, ST9, ST101, ST121, ST173, ST224, ST296, ST451, ST504) or NL only (ST18, ST37, ST54) (Fig. 3D). ST1 patients were significantly younger than all other listeriosis patients (*P*=3.5x10^-5^, *t*-test with Bonferroni-Holm correction). The higher number of MFL cases (that are of younger age) among the ST1 infections explains this effect, since no differences were observed when MFL cases were excluded. Differences in age distribution were not found for any of the other STs. At the serogroup level, infections with serogroup IIa strains were associated with reduced MFL and NL risks, while an above average MFL risk was detected for IVb infections (Fig. S2), to which hypervirulent ST1 belongs (Fig. 3D).

Information of patient sex was available for 1,550 isolates (86% of all isolates), according to which 42% were isolated from female patients (n=652) and 58% were from men (n=898). When MFL were excluded, general differences in sex distribution among patients infected with the 29 most prevalent STs shown in Fig. 3A were not detected, but a significant overrepresentation of female patients among all ST37 infections was observed (61% female, 39% male, *P*<0.05, χ^2^-test).

### *In vitro* strain virulence corresponds to risk potential of hyper- and hypovirulent STs

We selected one representative isolate per hyper- and hypovirulent ST identified above to test them in cell culture infection experiments. This selection included strain 18-04540 from the large Epsilon1a outbreak (18) as a representative for ST6 with average MFL/NL risk (Fig. 3D), strain 21-03201 from the Alpha10 cluster representing hypervirulent ST1, strain 19-05816 from the Pi4 cluster (40) for hypovirulent ST8, strain 19-06323 (Chi1a cluster) as a hypovirulent ST14 isolate, the sporadic 21-04322 strain representing hypovirulent ST29 and the Omikron1 strain 17-01049 as a member of hypovirulent ST155.

First, intracellular multiplication of these strains was determined in J774 mouse macrophages, but differences in uptake or intracellular replication were not observed (data not shown). Likewise, all strains formed plaques in 3T3 mouse fibroblasts indicating normal cell-to-cell spread (not shown). However, when the same experiment was repeated with HepG2 human liver cells, differences became apparent: For hypervirulent ST1 we generally observed a slightly increased invasion into HepG2 cells (compared to the reference ST6 isolate) in all experiments (1.5±0.3 fold in the experiment shown in Fig. 4). This was consistently found in all experiments but only reached statistical significance in 3 out of 4 repetitions. In contrast, invasion efficiency of hypovirulent ST8 (12±4% of the ST6 invasion level), ST14 (0.8±0.5%) and ST155 strains (6±0.3%) was significantly reduced. Consequently, final bacterial loads 6 hours post infection were also reduced to 22±10% (ST8), 0.9±0.2% (ST14) and 2.9±0.6 % of the ST6 reference level in the hypovirulent STs. In contrast, the ST29 isolate did not reveal an invasion or replication defect (Fig. 4) as suggested by its reduced disease severity (Fig. 3D). This shows that the ability to invade hepatocytes corresponds to the risk potential of the different STs to establish MFL and NL and is therefore of key importance for disease severity.

**Figure 4:**
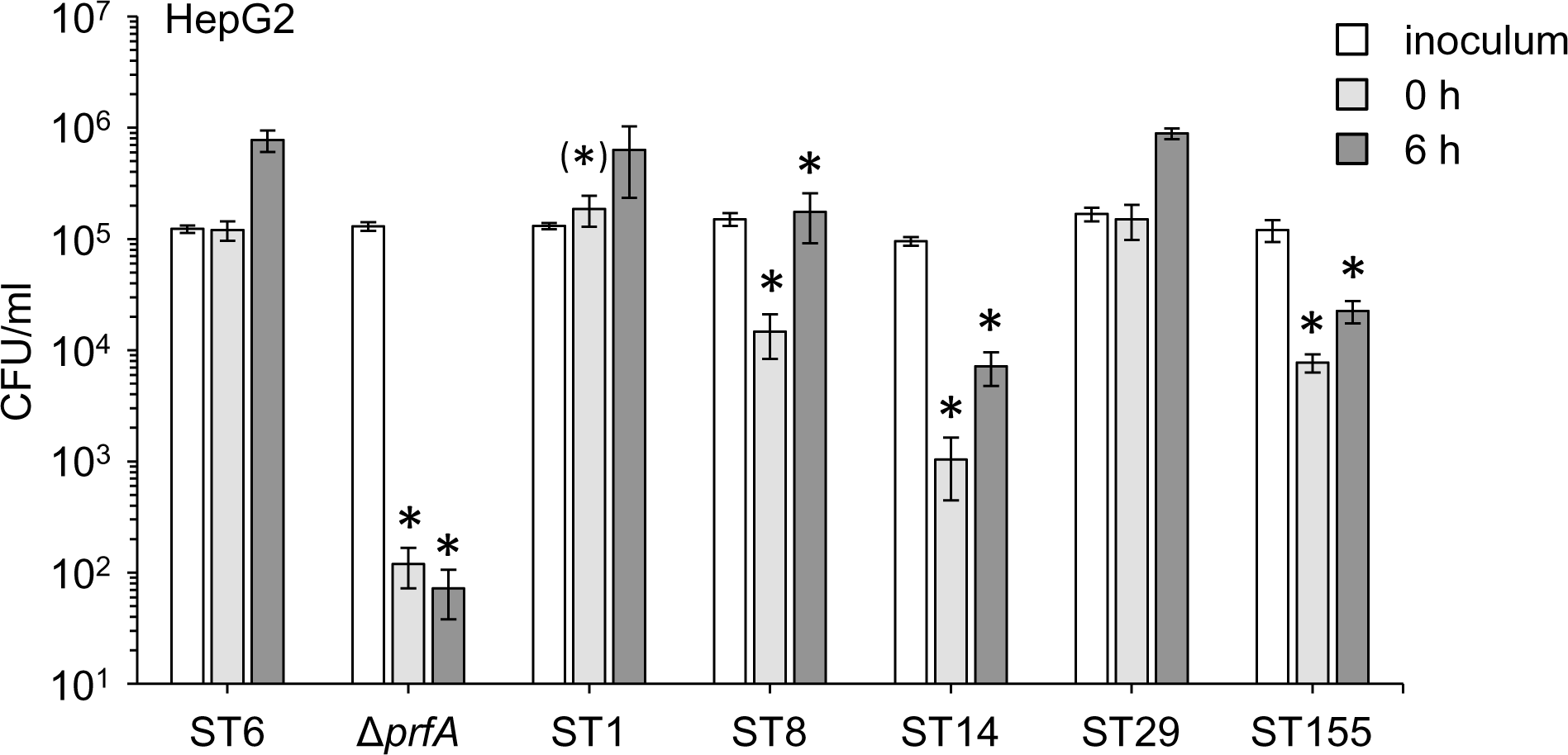
*In vitro* virulence of hyper- and hypovirulent STs. Invasion of and intracellular replication in HepG2 hepatocytes by representative *L. monocytogenes* strains belonging to hyper- and hypovirulent STs identified in this study. Strains tested were 18-04540 (ST6, Epsilon1a), 21-03201 (ST1, Alpha10), 19-05816 (ST8, Pi4), 19-06323 (ST14, Chi1a), 21-04322 (ST29, sporadic isolate) and 17-01049 (ST155, Omikron1). The experiment was carried out as triplicate and average values and standard deviations were calculated. Asterisks mark statistically significant differences compared to ST6 (*t*-test with Bonferroni-Holm correction, *P*<0.05).

### Size, duration and geographic distribution of listeriosis clusters

For determination of disease clusters, the 1,802 genomes were consecutively analysed by cgMLST using an initial threshold of ≤10 allele differences for cluster definition. For communication of a large number of outbreaks within the public/veterinary health offices, cgMLST clusters were named in the sequence of their detection using a nomenclature combining a Greek letter with a continuous number (*i. e.* Alpha1, Beta1,…Omega1, Alpha2, Beta2, and so on). Since March 2020, the cgMLST cluster cut-off was set to ≤7 allele differences, since larger distances were not observed in epidemiologically confirmed outbreaks (18, 30–32, 41). Clusters that could be differentiated into subclusters using the new threshold were labelled by additional letters (e. g. Epsilon1a, Epsilon1b,…). Over the four years, this approach grouped 1,129 of the 1,802 isolates (63%) into a cgMLST cluster and classified 673 isolates (37%) as sporadic (Fig. 2). As a result, isolates belonging to 253 different cgMLST clusters are included in the 2018-2021 isolate collection. For 65 of these clusters only one isolate fell into the study period, the remaining 188 clusters included 2-132 isolates (median: 3). The five largest clusters were Epsilon1a (132 isolates) (18), Ny9 (61 isolates) (42), Pi4 (51 isolates), Alpha10 (38 isolates) and Sigma1 (32 isolates) (31) (Fig. 2, Tab. 1).

**Tab. 1:**
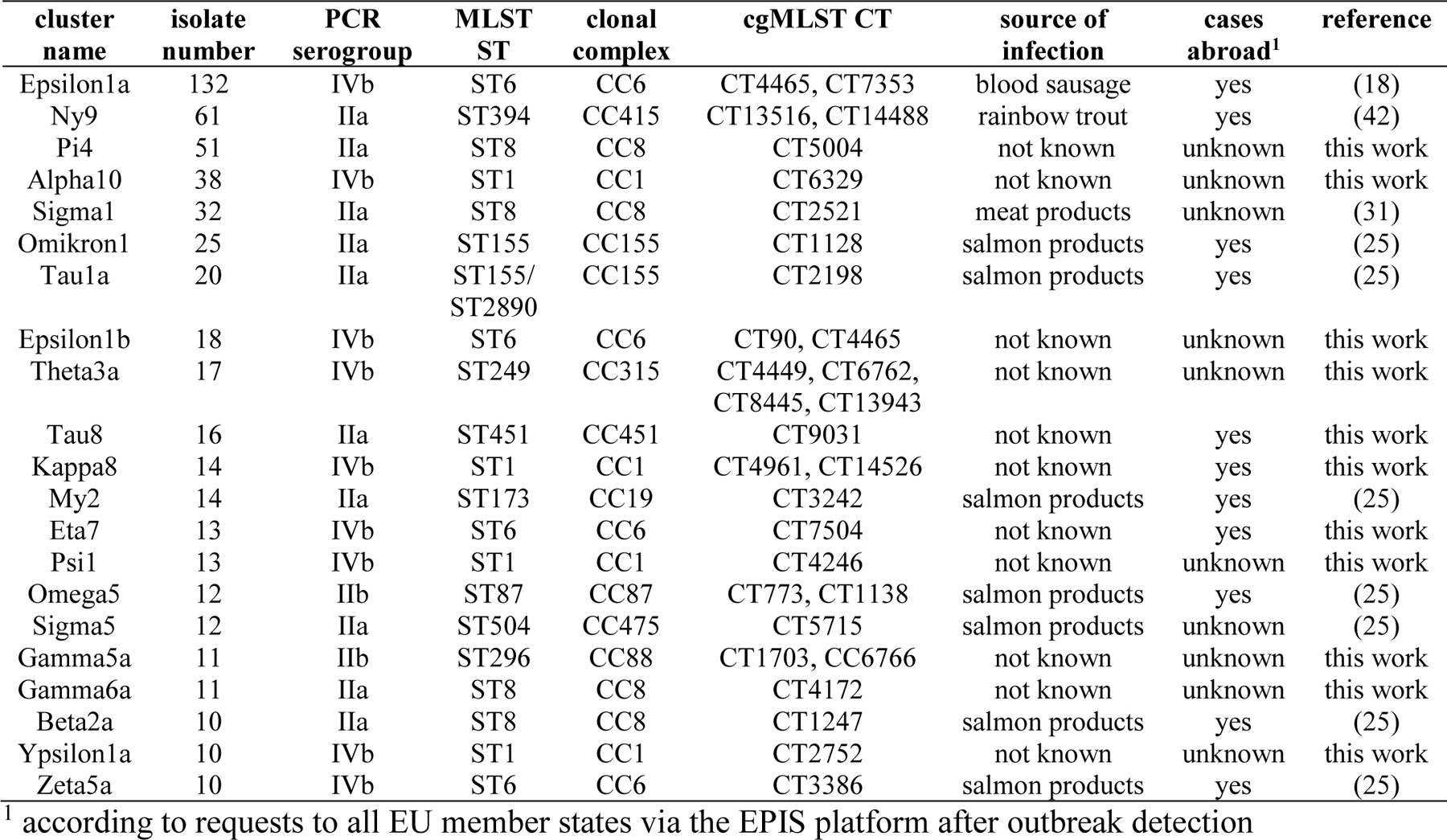
Key characteristics of German listeriosis clusters with ≥10 isolates in 2018-2021.

We defined cgMLST clusters as acute when all isolates belonging to this cluster were collected within 12 months and all remaining clusters as protracted. This grouped the 188 cgMLST clusters, into 109 acute and 79 protracted clusters (Fig. 4). The average period of activity of all these clusters together was 1.1 years (median: 0.7 years).

To determine their geographic distribution, we counted the number of federal countries for each cluster. Out of the 188 clusters, 75 clusters (40%) had cases in only one of the 16 German federal states and included 2-13 isolates (median: 2), we consider them as regional clusters. The remaining 113 clusters (60%) were considered cross-regional clusters, affected 2-13 federal states (median: 3) and included a higher number of isolates: 2-132 (median 4). Thus, a large proportion of the listeriosis outbreaks was active for more than one year and generated cases on a cross-regional scale. Clinical isolates matching several of these clusters from other European countries were identified through requests through the Epidemiological Intelligence System (EPIS) platform of the European Centre for Disease Control (ECDC), indicating that several outbreaks even generated cases on the European scale (Tab. 1).

### Significant German listeriosis clusters in 2018-2021

Twenty-one cgMLST clusters included ≥10 isolates out of which several clusters (Epsilon1a, Ny9, Sigma1) had been traced back to their infection sources and stopped (18, 31, 42) (Tab. 1). Seven more clusters with ≥10 isolates were linked to salmon consumption (25) (Tab. 1). However, several other large clusters (Pi4, Alpha10, Epsilon1b, Theta3a, Kappa8, Eta7) could not be traced back to their source yet (Tab.1).

**Pi4** comprised 51 isolates between 2018-2021 forming a contiguous cluster that differed in 0-16 cgMLST alleles from each other (Fig. 5A). Pi4 is geographically widespread in Germany, protracted (Fig. 5B-C) and closely related to but distinct from the clone, which caused the German Sigma1 outbreak recently (12-23 different cgMLST alleles) (31).

**Figure 5:**
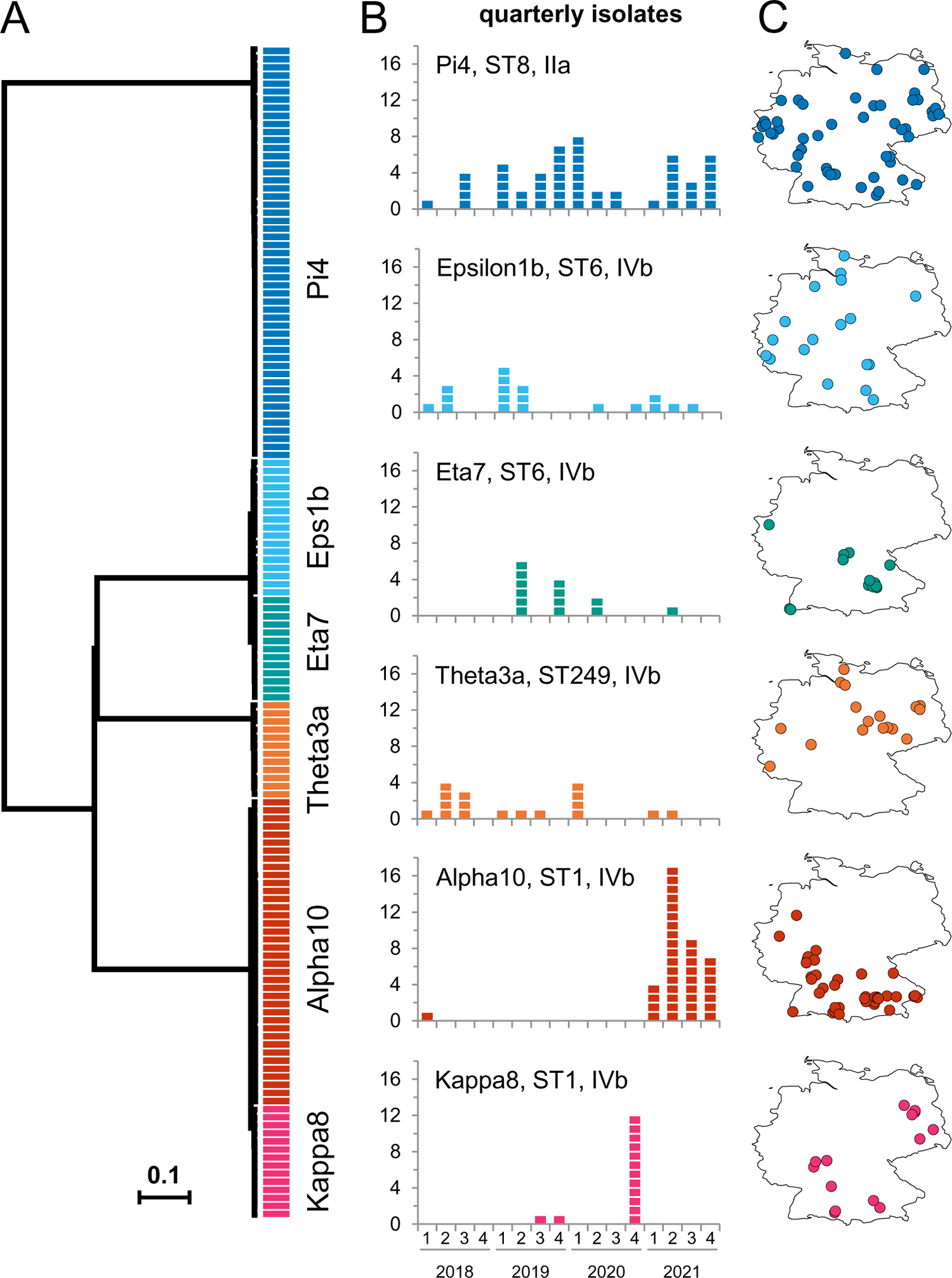
Large German listeriosis clusters with unknown infection source. (A) UPGMA tree calculated from 1,701 locus cgMLST data for the German listeriosis clusters Alpha10, Epsilon1b, Eta7, Kappa8, Pi4 and Theta3a. Isolates collected in 2018-2021 were included. (B) Epidemiological curves for the same clusters based on isolate collection dates. (C) Geographical origin of isolates within Germany.

**Alpha10** consists of 38 isolates (Tab. 1) and is highly clonal (0-3 alleles difference). Reconstruction of the closed genome of a representative Alpha10 isolate (21-03201, Tab. S2) and SNP calling showed that the Alpha10 isolates differed in 0-3 SNPs only. The majority of the Alpha10 isolates was collected in 2021 in South-Western Germany (Fig. 5B-C).

**Epsilon1b** includes 18 isolates (Tab. 1). In agreement with its protracted character (Fig. 5B), Epsilon1b is more heterogeneous and its isolates are distinguished from each other by 0-17 cgMLST alleles and 0-21 SNPs, with the reconstructed genome of Epsilon1b strain 11-04869 (Tab. S2) used as the reference. The Epsilon1b clone can be differentiated from the Epsilon1a clone, which caused the largest outbreak of invasive listeriosis detected in Germany so far (18), by the presence of a 42 kb prophage at the tRNA^Lys^ locus, which was collectively absent from the Epsilon1a isolates (Fig. S4).

**Theta3a** represents another protracted cluster and contains 17 isolates (Fig. 5B). Theta3a isolates belong to ST249, which are overrepresented in Germany (see above) and were collected from the North-Eastern and Western federal countries. They differ in 0-19 cgMLST alleles from each other and by 2-25 SNPs after variant calling using a reconstructed Theta3a genome (16-02236, Tab. S2) as the reference. Theta3a (ST249) constitutes a deeply branching clade within serogroup IVb. Interestingly, all ST249 strains carried an internal stop codon in *ispG* (ORY89_07620, *lmo1441* in EGD-e) (Tab. S3).

The **Kappa8** reflects a rather acute epidemiological incident with 14 isolates from Southern and Eastern Germany (Fig. 5B-C) that differed in 0-7 cgMLST alleles (median: 2) and 0-7 SNPs (median: 2) with the reconstructed genome of Kappa8 strain 19-07394 (Tab. S2) as the reference. **Eta7** is a cluster with pathogen isolation in West and South Germany (Fig. 5B). The 13 Eta7 isolates differed in 0-5 (median: 2) cgMLST alleles and 0-9 SNPs (median: 7) using the genome of Eta7 strain 19-02390 as the reference (Tab. S2). Remarkably, 12 of the 13 Eta7 isolates were collected from new-borns or pregnant women.

### Maintenance of virulence and fitness gene function in hypervirulent IVb isolates

To explain reduced MFL and NL risks of hypovirulent STs, we inspected the LIPI-1 and *inl* internalin genes for the presence of premature stop codons (PMSCs) and frameshifts. Inactivating mutations were not found in the LIPI-1 and *inlB, inlC, inlE, inlGH, inlI, inlJ, inlL and inlP* genes in any of the clinical isolates, but were present in *inlA* of all ST121 isolates and in several ST9 clones (Tab. S3). Moreover, all Tau8 strains (ST451) carried a PSMC in the *inlF* gene (Tab. S3). InlF was reported to support the uptake of *L. monocytogenes* by mouse macrophages (43). However, the *inlF*⁻ Tau8 isolate 20-01331 was taken up and replicated to the same bacterial titer in J774 mouse macrophages as efficiently as the EGD-e wild type or two other InlF positive ST451 strains belonging to the Xi5 and Omikron5 clusters (Fig. 6A).

**Figure 6:**
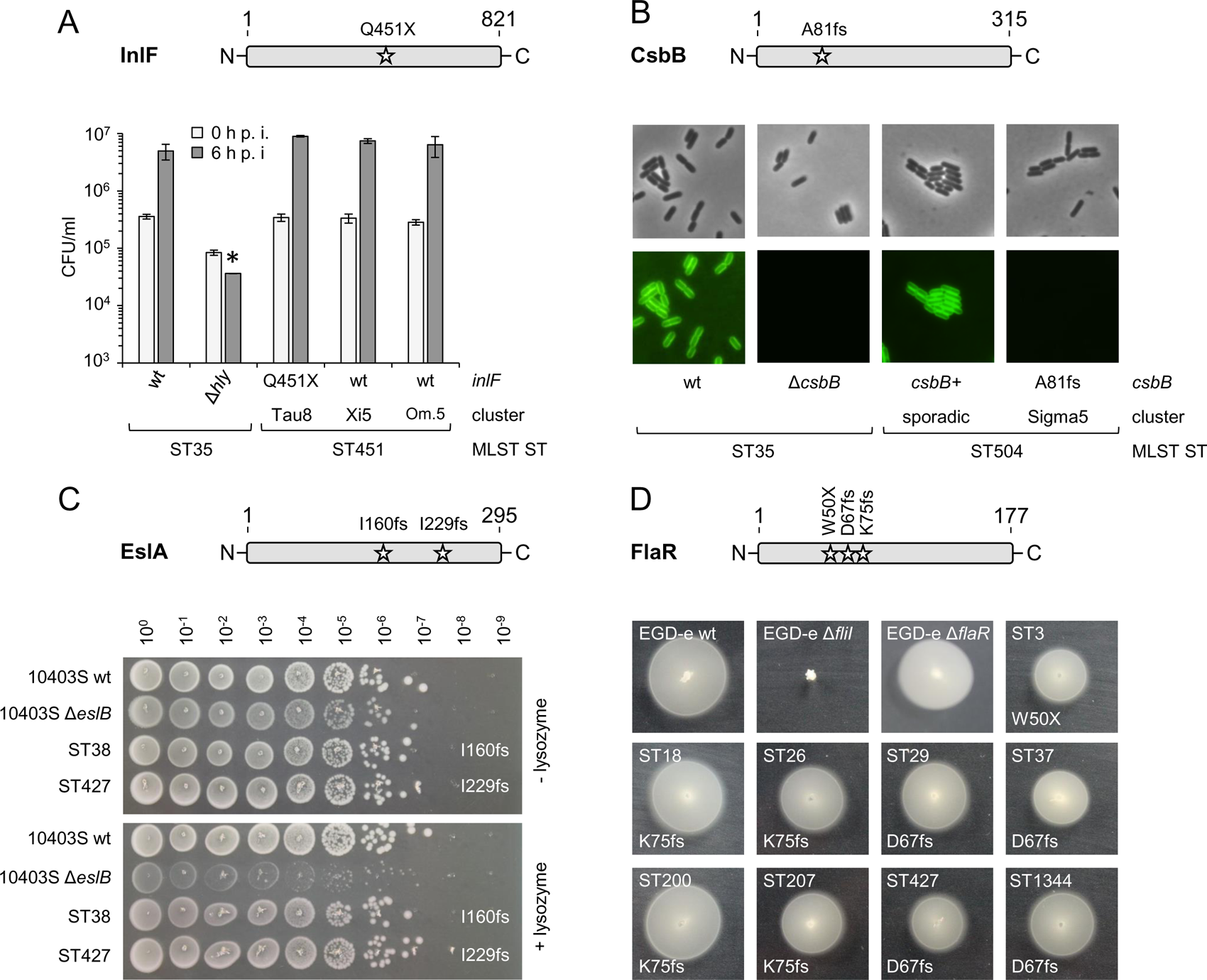
Phenotypes of clinical isolates with inactivated *inlF*, *csbB*, *eslA* and *flaR* genes. (A) A premature stop codon inactivates the *inlF* gene of Tau8 isolates (ST451, CT9031). Scheme showing the position of the Q451X mutation within *inlF* of Tau8 strains (upper panel). Despite inactivated *inlF*, The Tau8 isolate 20-01331 shows full virulence in a mouse macrophage infection assay (lower panel). J774 mouse macrophage were infected with *L. monocytogenes strains* EGD-e (ST35, wt), LMS250 (ST35, Δ*hly*), 20-01331 (Tau8, ST451, InlF truncated), 18-03445 (Xi5, ST451, full-length InlF) and 18-02122 (Omikron5, ST451, full-length InlF) and the bacterial titer was determined right after infection (0 h p. i.) and 6 hours later (6 h p. i.). Average values and standard deviations were calculated from technical triplicates. The asterisk indicates a statistically significant difference (*P*<0.01, *t*-test with Bonferroni-Holm correction). Abbr.: Om.5 – Omikron5. (B) Sigma5 isolates (ST504, CT5715) carry an inactivated *csbB* gene. Scheme illustrating the position of the A81fs mutation within *csbB* of Sigma5 strains (upper panel). Micrographs of *L. monocytogenes* strains 20-06257 (ST504, CT14635, sporadic, *csbB*^+^) and 21-06873 (ST504, CT5715, Sigma5, *csbB* A81fs) after staining with fluorescently labelled wheat germ agglutinin (lower panel). Phase contrast (top row) and fluorescence images (bottom row) are shown. ST35 strains EGD-e (wt) and LMJR156 (Δ*csbB*) were included as controls. (C) ST38 and ST427 isolates carry *eslA* inactivating mutations. Scheme showing the position of the inactivating mutations within *eslA* (upper panel). Spot dilution assay to determine lysozyme resistance of representative ST38 (20-02710) and ST427 (21-00930) isolates on BHI agar plates ± lysozyme (lower panel). ST87 strains 10403S (wt) and ANG4275 (10403S Δ*eslB*) mutant were included as controls. (D) Mutations inactivating the *flaR* gene in various subtypes. Scheme showing the position of *flaR* inactivating mutations (upper panel). Swarming assay to test flagellar motility of representative ST3 (18-04580), ST18 (18-00242), ST26 (19-02197), ST29 (18-03980), ST37 (20-01921), ST200 (18-02068), ST207 (20-01871), ST427 (18-01591) and ST1344 (21-01230) isolates (lower panel). Isogenic strains EGD-e, LMSW211 (Δ*flaR*) and LMS3 (Δ*fliI*, all belonging to ST35) were included as controls. Respective STs and their *flaR* mutations are indicated.

Comparison of cgMLST/agMLST allele numbers with a previously generated list of inactivated *L. monocytogenes* alleles (44) further detected inactivating mutations in the autolysin encoding *aut* gene in several clones belonging to the ST3, ST121 and hypovirulent ST155 sublineages and an inactivating mutation in the *chiB* chitinase gene in several ST4 isolates (Tab. S3). PSMCs were also identified in the chitin binding protein encoding *lmo2467* gene in the majority of the hypovirulent ST29 isolates and in the My5 cluster belonging to ST9 (Tab. S3).

The *csbB* gene, required for decoration of wall teichoic acids with N-acetylglucosamine (GlcNAc) (45), carried frameshift mutations in clones belonging to ST87 and ST504. Fluorescent lectin staining to detect the presence of GlcNAc-modified wall teichoic acids (WTA) showed that WTAs of the *csbB*⁻ Sigma5 (ST504) isolate 21-06873 were not decorated with GlcNAc as expected, whereas no such defect was observed in a related sporadic *csbB^+^* ST504 isolate (Fig. 6B). We also found that the *eslA* gene, encoding the ATPase of the EslABC transporter, which contributes to lysozyme resistance (46), was frameshifted in all ST38 and ST427 isolates (Tab. S3). However, two selected ST38 and ST247 strains showed the same resistance to lysozyme as wild type strain 10403S in contrast to its isogenic Δ*eslB* mutant (Fig. 6C).

The *clpP1* protease gene of all nine ST16 strains and the *ispG* gene from the non-mevalonate pathway of all ST249 strains was also truncated. ST16 belongs to the same clonal complex as hypovirulent ST8, while ST249 is a sublineage within serogroup IVb, but one with an average disease severity scores (Fig. 3D). The motility gene *flaR* was truncated in most ST3 isolates and frameshifted in most if not all ST18, ST26, hypovirulent ST29, ST37, ST200, ST207, ST427 and ST1344 strains (Tab. S3). However, motility of representative isolates from these *flaR*⁻ STs and of a Δ*flaR* deletion mutant was not impaired (Fig. 6D).

Remarkably, 23% of the IIa (hypovirulent), half of the IIc (reduced MFL risk) and 20% of the IIb isolates (average MFL/NL risk) carried one of the mutations identified above, but only 4% of the IVb isolates (hypervirulent), and out of these only ST249 (*ispG*) as well as some ST4 isolates (*chiB*) were affected. (Tab. S3). Thus, it seems that purifying selection maintains the function of these genes in the majority of the IVb isolates. Moreover, the importance of these genes for *L. monocytogenes* biology originally established in reference strains cannot be generalized.

## Discussion

### Population structure and disease clusters

The combination of sequencing data covering the majority of clinical cases with notification data enabled us to analyse human *L. monocytogenes* infections that occurred in Germany over a period of four consecutive years in different dimensions. First, we observed the same bipartite population structure as described by others (47, 48) with two genetically distinct lineages that each contributed to almost equal parts to listeriosis cases in Germany: Phylogenetic lineage I (comprising serogroups IIb, IVb, IVbv-1) and phylogenetic lineage II (IIa and IIc), whereas strains of other lineages were only found rarely or not at all. Separation into two well-separated lineages likely reflects pathogen adaptation to different reservoirs. Second, the overall population structure did not change much compared to a previous analysis covering isolates from 2007-2017 (20). During both periods, serogroups IVb, IIa and IIb as well as sequence types ST6, ST1, ST8 were the three most prevalent phylogenetic groups in Germany (20). Stable colonization of environmental habitats and infection sources with established and persistent clones, respectively, and rarely occurring infiltration of these niches by new subtypes from external ecosystems would explain this observation (49). Furthermore, the majority of isolates belonged to cgMLST clusters, out of which more than the half of the clusters generated cases on a cross-regional scale. Thus, foodstuff produced for supra-regional retail rather than local production and consumption accounts for most listeriosis cases in Germany. A significant portion of the listeriosis clusters also is active for more than 12 months, which would be consistent with (i) constant pathogen dissemination from environmental sources into the food processing chain, (ii) persistent contamination of food processing facilities with the same clone or (iii) long-term storage of contaminated products in patient households. At least for the latter two scenarios, several published examples exist that made them appear plausible (30, 50, 51). Several of the clusters identified also reflected real listeriosis outbreaks (Tab.1). Their recognition can initiate successful back-tracing of *L. monocytogenes* clones to their infection source as demonstrated during past outbreaks (17–19).

### Risk profiling of prevalent clinical *L. monocytogenes* subtypes

Besides its importance for public health, genomic pathogen surveillance also allows the identification of hypo- and hypervirulent subtypes or regionally prevailing clones, when genomic subtyping data are combined with information on disease severity or geographic data, respectively. Using this type of approach, the second most prevalent subtype (ST1) turned out as hypervirulent when compared to ST6 showing average risk potential for development of MFL/NL, while others (ST8, ST14, ST29 and ST155) generated less MFL and NL cases. Remarkably, ST1 strains had also been associated with increased rates of MFL/NL infections by other authors (26, 52), and classification of ST8 and ST155 strains as less virulent further validates our data, as this is consistent with the reduced rates of MFL/NL infections or their experimentally proven hypovirulence, respectively, reported in other studies (26, 53). Furthermore, hypovirulence of ST14 isolates seen here is in accordance with their reduced *in vitro* invasion efficiency into Caco-2 cells (54) and their previous classification as an environment-associated subtype (55). However, this view is challenged by reports showing increased virulence in *Galleria* infection assays of CC14 (56). ST29 represents a newly discovered subtype associated with reduced MFL/NL rates. This is congruent with the detection of ongoing gene loss (*flaR*, *lmo2467*) in this phylogenetic clade (this work), probably resulting in fitness defects at least under certain conditions. The observation that several STs were associated with decreased risks for MFL or NL without affecting the risk for the respective other disease condition is consistent with the idea that pathogen transmission to the brain and the placenta is supported by specific sets of virulence factors, the functionality of which can be impaired independently from each other. However, inactivation of the known determinants for brain (*inlF*) (57) or placenta invasion (*inlP*) (58) did not explain the occurrence of several STs with reduced risk for either MFL or NL. We also did not detect a significantly reduced NL risk associated with Tau8 infections (truncated *inlF*), but the small number of Tau8 patients, for which clinical data were available, has probably hampered detection of a significant effect. Significant associations were found between disease manifestations and serogroups, while presence, absence or total number of pathogenicity islands were not associated with disease forms. Quantification of such associations in our approach is surely masked by non-genetic confounders such as consumer behaviour, possible biases in the differential consumption of outbreak-associated food items in certain risk groups or other patient-associated influences. As further limitations, data on disease manifestation was not available for all isolates/cases and data on the immune status or comorbidities were not generally available. Despite these limitations, our study represents one of the very few surveys currently available (26) that combines genomic pathogen typing data with clinical data on disease outcome from a systematic national sampling program that almost achieves full case coverage. Our analysis also suggests that further genetic factors determining infection of the brain and the placenta may exist, as secondary organ involvement has to be considered a multifactorial process (52).

### Further benefits of genomic surveillance systems

As another benefit, genomic pathogen surveillance allows the measurement of the frequency of natural gene loss that occurs in clinical pathogen populations. Since the majority of our isolates were collected from invasive listeriosis, *i. e.* isolated from primary sterile body fluids, genes found to be inactivated by PSMCs or frameshift mutations in a sufficient number of isolates (*e. g. csbB*, *clpP1*, *eslA*, *flaR*, *inlA* or *inlF*) are likely not essential for pathogen transmission from the gut to the blood stream *per se*. In this way, genomic pathogen surveillance provides an unbiased possibility to verify or relativize data on the importance of selected genes for *L. monocytogenes* virulence that had been deduced from *in vitro* or animal studies. For example, even though ClpP1 is secondary to ClpP2, and only supports ClpP2 function (59), we were actually surprised to see that *clpP1* can be lost in clinical isolates having caused invasive disease, since ClpP proteins have crucial roles in protein homeostasis (60). Apparently, *clpP1* likely is an accessory or even remnant gene that is not essential for survival in the environment or during systemic human infection. Thus, analysis of the data generated by genomic surveillance systems permits conclusions on the relevance of genes for pathogen biology, ultimately supporting functional annotation of the *L. monocytogenes* genome (61). Besides this, genomic pathogen surveillance exerted on the global scale helps to reconstruct pathogen transmission routes across countries and allows identification of globally disseminated as well as regionally dominating subtypes (49). According to our results, the majority of the STs that caused disease in Germany has also been reported from locations abroad. However, numerous subtypes exist that have so far only been found outside Germany while others (e. g. ST173 and ST249) were overrepresented in Germany at the time of analysis. Thus, the population of *L. monocytogenes* subtypes that is pathogenic to humans is composed of internationally widespread clones and subtypes with regionally restricted distribution.

Taken together, this data set combined with our conclusions provides a comprehensive insight into the population structure of clinical *L. monocytogenes* isolates in Germany and the genetic and clinical characteristics of the most abundant phylogenetic subtypes. The assignment of hyper- and hypovirulent lineages may help to prioritize clusters for epidemiological investigations and to concentrate future work on the identification of the underlying genetic determinants, which might lead to the discovery of relevant virulence factors outside of reference strains.

## Methods

### Bacterial strains and growth conditions

All clinical strains used in this study are listed in Tab. S1. Samples were accompanied by sample submission forms that included basic information on the source of isolation as well as the disease manifestation. *L. monocytogenes* strains were routinely grown in brain heart infusion (BHI) broth or on BHI agar plates at 37°C. *L. monocytogenes* strains EGD-e (wild type IIc strain, ST35, CC9) (62) and its isogenic descendants BUG2214 (Δ*prfA*) (63), LMS3 (Δ*fliI*) (64), LMS250 (Δ*hly*) (44), LMJR156 (Δ*csbB*) (65), LMSW211 (Δ*flaR*, this work) as well as strain 10403S (wild type IIa strain, ST87, CC7) and its isogenic Δ*eslB* mutant (46) were included as controls in selected experiments.

### Whole genome sequencing, PCR serogrouping, MLST and cgMLST

DNA was isolated by mechanical disruption using glass beads in a TissueLyser II bead mill (Qiagen, Hilden, Germany) (66) and quantified with a Qubit dsDNA BR (or HS) Assay kit and Qubit fluorometers (Invitrogen, Carlsbad, CA, USA). Libraries were prepared using the Nextera XT DNA Library Prep Kit and sequenced on MiSeq or NextSeq sequencers in 1 x 150 bp single end or 2 x 250 bp or 2 x 300 bp paired end mode. A SeqSphere (Ridom, Münster, Germany) script was used for read trimming and contig assembly with Velvet as the assembler. *In silico* PCR serogroups, seven locus multi-locus sequence typing (MLST) sequence types (STs) and 1,701 locus core genome MLST (cgMLST) complex types (CTs) were automatically extracted using SeqSphere (67). Likewise, cgMLST clusters and minimum spanning trees were calculated in SeqSphere in the “pairwise ignore missing values” mode and annotated using iTOL (68).

### Matching of isolates with notified cases and calculation of risk differences

According to the German Protection Against Infection Act, laboratory confirmation of *L. monocytogens* isolation or detection of nucleic acids from blood, cerebrospinal fluid, or other usually sterile sites is notifiable to local health authorities and are electronically transmitted to the RKI. Case notification through the German notification system partially includes information on disease manifestation (listeriosis of neonates, listeriosis of pregnant women, other forms of listeriosis) and symptoms (meningitis, sepsis among others). These categories were combined with isolate identifiers and typing information through merging of isolates with notification cases.

Confirmed cases of materno-fetal infections (MFL) and neurolisteriosis (NL) were counted per ST group and risk differences and 95% confidence intervals were calculated for each individual ST compared to all other STs according to standard procedures (69).

### Generation of closed genomes

For the generation of closed genome sequences, DNA of the bacterial strains was extracted with the GenEluteTM Bacterial Genomic DNA Kit (Sigma). The libraries were prepared with the Rapid barcoding kit (SQK-RBK004, Oxford Nanopore Technologies, Oxford, UK) kit and afterwards subjected to sequencing in a FLOMIN 106D flow-cell on a MinION device (Oxford Nanopore Technologies, Oxford, UK). The obtained long reads were filtered using the Filtlong tool (70) with standard settings and the following modifications: the minimal length of reads was set to 3,000 bp and the target bases to 290,000,000 bp (100-fold coverage). For filtering, the corresponding Illumina reads were used as an external reference. The improved subset of long reads and the Illumina reads were used in a hybrid assembly with Unicycler (71) to generate a closed genome sequence. Genomes were annotated using the NCBI prokaryotic genome annotation pipeline.

### Single nucleotide polymorphism-based alignments and whole genome MLST

The batchMap pipeline (20) was used for mapping of sequencing reads against closed reference genomes, for generation of consensus sequences and alignments. Single nucleotide polymorphisms (SNP) were filtered using the SNPfilter pipeline (72) and an exclusion distance of 300 nt. Phylogenetic trees were calculated using SeqSphere or Geneious (Biomatters Ltd., Auckland, New Zealand) and annotated in iTOL (68). For whole genome MLST (wgMLST) analysis, non-redundant open reading frames from closed genomes of selected strains were automatically extracted in SeqSphere and used for generation of an *ad hoc* wgMLST schemes in SeqSphere. Allele calling and analysis procedures were the same as for standard cgMLST.

### Allele calling for virulome and resistome analysis

Known *L. monocytogenes* virulence and resistance genes were included as target loci in a SeqSphere task template (18). Assembled genomes were queried against these task templates in SeqSphere for the presence or absence of these genes using an identity cut-off of 90% and an alignment coverage cut-off of 99%.

### Identification of premature stop codons

We compared the cgMLST and accessory genome MLST (agMLST) allele annotations of the 1,802 isolates with a previously generated list of *L. monocytogenes* cg/agMLST alleles affected by internal stop codons (44). As the cg/agMLST algorithm annotates frameshifted alleles as “failed” alleles, we also inspected the cg/agMLST alleles for the occurrence of failed alleles that are associated with particular sequence types or even specific cgMLST clusters. We considered a gene as inactivated, when an allele variation generated a stop codon between the first 5% and the last 80% of an open reading frame sequence and further included only those genes in the analysis that were affected at least twice within the same phylogroup to exclude accidental sequencing errors.

### Construction of a Δ*flaR* mutant

To remove *flaR* (*lmo1412*) from the chromosome, regions up- and downstream of *flaR* were amplified using the oligonucleotides SW200 (GATCTATCGATGCATGCCATGGCGATTAGTTCTGTTATAATGGTTATTAGC)/ SW202 (GCTATTTATCACATTTTAAGCACTCCTTATCTGACTATG) and SW203 (GCTTAAAATGTGATAAATAGCCCATGAATGCTTGG)/ SW214 (GCGCGCGTCGACCAAGTACCATCAAATCAATCCGGAAC), respectively, as primers and fused together by splicing by overlapping extension PCR. The Δ*flaR* fragment was inserted into pHoss1 (73) using NcoI/SalI, and the resulting plasmid (pSW102) was transferred to *L. monocytogenes* EGD-e by electroporation. The *flaR* gene was then deleted following the plasmid insertion/excision protocol of Abdelhamed *et al.* (73). Removal of *flaR* and loss of pSW102 was confirmed by PCR and the resulting strain was named LMSW211.

### Infection experiments

Infection of J774 mouse macrophages, HepG2 cells and 3T3 fibroblasts with *L. monocytogenes* strains was performed as described earlier (41).

### Phenotypic assays

For determination of lysozyme sensitivity, *L. monocytogenes* strains were grown over night in BHI broth at 37°C and aliquots of a ten-fold dilution series were spotted on BHI agar plates ± 100 µg/ml lysozyme. Agar plates were incubated over night at 37°C and then photographed.

For analysis of flagellar motility, strains grown on BHI agar plates were stab-inoculated into LB agar plates containing 0.3% (w/v) agarose. The plates were incubated over night at 30°C and photographed the next morning.

In order to visualize wall teichoic acid decoration with N-acetylglucosamine, *L. monocytogenes* cells taken from overnight cultures were diluted 1:100 in fresh BHI and grown for 4 hours at 37°C. Cells from 100 µl aliquots were collected by centrifugation and washed twice with PBS buffer. The washed cells were stained with 0.1 mg/ml CF^®^488A wheat germ agglutinin conjugate (Biotium, Fremont, CA, USA) in PBS for 5 min at room temperature and washed two more times with PBS buffer. An 0.5 µl aliquot of the stained cell suspension was spotted onto a microscope slide covered with a thin film of 1.5% agarose and covered with a cover slip. Images were taken using a Nikon Eclipse Ti microscope coupled to a Nikon Mono DS-Qi2 CMOS camera and processed using the NIS elements AR software package (Nikon).

### Statistics

Levels of significance were determined using a standard two-tailed *t*-test and weighted using the Bonferroni-Holm correction. Values below the threshold of *P*<0.05 were considered as statistically significant.

## Data availability

Genome sequencing data are available at the [European Nucleotide Archive](https://www.ebi.ac.uk/ena/browser/home) using the accession numbers given in supplementary table S1.

## Acknowledgments

We thank Simone Dumschat and Birgitt Hahn for excellent technical assistance and the Genome Competence Centre of the RKI for sequencing of *L. monocytogenes* genomes. We also would like to thank all the primary diagnostic labs for submission of *L. monocytogenes* strains. We further acknowledge Jeanine Rismondo (University of Göttingen) and Angelika Gründling (Imperial College, London) for sharing strain 10403S and the Δ*eslB* mutant. This work was funded by DFG grant HA6830/5-1 (to SH) and financing by the German Ministry of Health dedicated to the Consultant Laboratory for *Listeria* (to AF). The funders had no role in the design of the study, collection and analysis of data, decision to publish or preparation of the manuscript.

## References

1. Matle I, Mbatha KR, Madoroba E. 2020. A review of *Listeria monocytogenes* from meat and meat products: Epidemiology, virulence factors, antimicrobial resistance and diagnosis. Onderstepoort J Vet Res 87:e1–e20.

2. Wilking H, Lachmann R, Holzer A, Halbedel S, Flieger A, Stark K. 2021. Ongoing High Incidence and Case-Fatality Rates for Invasive Listeriosis, Germany, 2010-2019. Emerg Infect Dis 27:2485–2488.

3. Pohl AM, Pouillot R, Bazaco MC, Wolpert BJ, Healy JM, Bruce BB, Laughlin ME, Hunter JC, Dunn JR, Hurd S, Rowlands JV, Saupe A, Vugia DJ, Van Doren JM. 2019. Differences Among Incidence Rates of Invasive Listeriosis in the U.S. FoodNet Population by Age, Sex, Race/Ethnicity, and Pregnancy Status, 2008-2016. Foodborne Pathog Dis 16:290–297.

4. 4. Authority EFS, Prevention ECfD, Control. 2021. The European Union One Health 2020 Zoonoses Report. EFSA Journal 19:e06971.

5. Vazquez-Boland JA, Kuhn M, Berche P, Chakraborty T, Dominguez-Bernal G, Goebel W, Gonzalez-Zorn B, Wehland J, Kreft J. 2001. *Listeria* pathogenesis and molecular virulence determinants. Clin Microbiol Rev 14:584–640.

6. Quereda JJ, Moron-Garcia A, Palacios-Gorba C, Dessaux C, Garcia-Del Portillo F, Pucciarelli MG, Ortega AD. 2021. Pathogenicity and virulence of *Listeria monocytogenes*: A trip from environmental to medical microbiology. Virulence 12:2509–2545.

7. Scallan E, Hoekstra RM, Angulo FJ, Tauxe RV, Widdowson MA, Roy SL, Jones JL, Griffin PM. 2011. Foodborne illness acquired in the United States--major pathogens. Emerg Infect Dis 17:7–15.

8. Charlier C, Perrodeau E, Leclercq A, Cazenave B, Pilmis B, Henry B, Lopes A, Maury MM, Moura A, Goffinet F, Dieye HB, Thouvenot P, Ungeheuer MN, Tourdjman M, Goulet V, de Valk H, Lortholary O, Ravaud P, Lecuit M, group Ms. 2017. Clinical features and prognostic factors of listeriosis: the MONALISA national prospective cohort study. Lancet Infect Dis 17:510–519.

9. Werber D, Hille K, Frank C, Dehnert M, Altmann D, Müller-Nordhorn J, Koch J, Stark K. 2013. Years of potential life lost for six major enteric pathogens, Germany, 2004-2008. Epidemiol Infect 141:961–8.

10. Autio T, Markkula A, Hellstrom S, Niskanen T, Lunden J, Korkeala H. 2004. Prevalence and genetic diversity of *Listeria monocytogenes* in the tonsils of pigs. J Food Prot 67:805–8.

11. Hellstrom S, Laukkanen R, Siekkinen KM, Ranta J, Maijala R, Korkeala H. 2010. *Listeria monocytogene*s contamination in pork can originate from farms. J Food Prot 73:641–8.

12. Walland J, Lauper J, Frey J, Imhof R, Stephan R, Seuberlich T, Oevermann A. 2015. *Listeria monocytogenes* infection in ruminants: Is there a link to the environment, food and human health? A review. Schweiz Arch Tierheilkd 157:319–28.

13. Vivant AL, Garmyn D, Piveteau P. 2013. *Listeria monocytogenes*, a down-to-earth pathogen. Front Cell Infect Microbiol 3:87.

14. Miceli A, Settanni L. 2019. Influence of agronomic practices and pre-harvest conditions on the attachment and development of *Listeria monocytogenes* in vegetables. Annals of Microbiology 69:185–199.

15. Mazaheri T, Cervantes-Huaman BRH, Bermudez-Capdevila M, Ripolles-Avila C, Rodriguez-Jerez JJ. 2021. *Listeria monocytogenes* Biofilms in the Food Industry: Is the Current Hygiene Program Sufficient to Combat the Persistence of the Pathogen? Microorganisms 9.

16. EU. 2020. Commission Regulation (EC) No 2073/2005 of 15 November 2005 on microbiological criteria for foodstuffs. http://data.europa.eu/eli/reg/2005/2073/2020-03-08.

17. Thomas J, Govender N, McCarthy KM, Erasmus LK, Doyle TJ, Allam M, Ismail A, Ramalwa N, Sekwadi P, Ntshoe G, Shonhiwa A, Essel V, Tau N, Smouse S, Ngomane HM, Disenyeng B, Page NA, Govender NP, Duse AG, Stewart R, Thomas T, Mahoney D, Tourdjman M, Disson O, Thouvenot P, Maury MM, Leclercq A, Lecuit M, Smith AM, Blumberg LH. 2020. Outbreak of Listeriosis in South Africa Associated with Processed Meat. N Engl J Med 382:632–643.

18. Halbedel S, Wilking H, Holzer A, Kleta S, Fischer MA, Luth S, Pietzka A, Huhulescu S, Lachmann R, Krings A, Ruppitsch W, Leclercq A, Kamphausen R, Meincke M, Wagner-Wiening C, Contzen M, Kraemer IB, Al Dahouk S, Allerberger F, Stark K, Flieger A. 2020. Large Nationwide Outbreak of Invasive Listeriosis Associated with Blood Sausage, Germany, 2018-2019. Emerg Infect Dis 26:1456–1464.

19. Fernandez-Martinez NF, Ruiz-Montero R, Briones E, Banos E, Garcia San Miguel Rodriguez-Alarcon L, Chaves JA, Abad R, Varela C, team L, Lorusso N. 2022. Listeriosis outbreak caused by contaminated stuffed pork, Andalusia, Spain, July to October 2019. Euro Surveill 27.

20. Halbedel S, Prager R, Fuchs S, Trost E, Werner G, Flieger A. 2018. Whole-Genome Sequencing of Recent *Listeria monocytogenes* Isolates from Germany Reveals Population Structure and Disease Clusters. J Clin Microbiol 56.

21. Moura A, Tourdjman M, Leclercq A, Hamelin E, Laurent E, Fredriksen N, Van Cauteren D, Bracq-Dieye H, Thouvenot P, Vales G, Tessaud-Rita N, Maury MM, Alexandru A, Criscuolo A, Quevillon E, Donguy MP, Enouf V, de Valk H, Brisse S, Lecuit M. 2017. Real-Time Whole-Genome Sequencing for Surveillance of *Listeria monocytogenes*, France. Emerg Infect Dis 23:1462–1470.

22. Kwong JC, Mercoulia K, Tomita T, Easton M, Li HY, Bulach DM, Stinear TP, Seemann T, Howden BP. 2016. Prospective Whole-Genome Sequencing Enhances National Surveillance of *Listeria monocytogenes*. J Clin Microbiol 54:333–42.

23. Van Walle I, Bjorkman JT, Cormican M, Dallman T, Mossong J, Moura A, Pietzka A, Ruppitsch W, Takkinen J, European Listeria Wgs Typing G. 2018. Retrospective validation of whole genome sequencing-enhanced surveillance of listeriosis in Europe, 2010 to 2015. Euro Surveill 23.

24. Jackson BR, Tarr C, Strain E, Jackson KA, Conrad A, Carleton H, Katz LS, Stroika S, Gould LH, Mody RK, Silk BJ, Beal J, Chen Y, Timme R, Doyle M, Fields A, Wise M, Tillman G, Defibaugh-Chavez S, Kucerova Z, Sabol A, Roache K, Trees E, Simmons M, Wasilenko J, Kubota K, Pouseele H, Klimke W, Besser J, Brown E, Allard M, Gerner-Smidt P. 2016. Implementation of Nationwide Real-time Whole-genome Sequencing to Enhance Listeriosis Outbreak Detection and Investigation. Clin Infect Dis 63:380–6.

25. Lachmann R, Halbedel S, Lüth S, Holzer A, Adler M, Pietzka A, Dahouk SA, Stark K, Flieger A, Kleta S, Wilking H. 2022. Invasive listeriosis outbreaks and salmon products: a genomic, epidemiological study. Emerg Microbes Infect doi:10.1080/22221751.2022.2063075:1-30.

26. Maury MM, Tsai YH, Charlier C, Touchon M, Chenal-Francisque V, Leclercq A, Criscuolo A, Gaultier C, Roussel S, Brisabois A, Disson O, Rocha EP, Brisse S, Lecuit M. 2016. Uncovering *Listeria monocytogenes* hypervirulence by harnessing its biodiversity. Nat Genet 48:308–13.

27. Harter E, Wagner EM, Zaiser A, Halecker S, Wagner M, Rychli K. 2017. Stress Survival Islet 2, Predominantly Present in *Listeria monocytogenes* Strains of Sequence Type 121, Is Involved in the Alkaline and Oxidative Stress Responses. Appl Environ Microbiol 83.

28. Kremer PH, Lees JA, Koopmans MM, Ferwerda B, Arends AW, Feller MM, Schipper K, Valls Seron M, van der Ende A, Brouwer MC, van de Beek D, Bentley SD. 2017. Benzalkonium tolerance genes and outcome in *Listeria monocytogenes* meningitis. Clin Microbiol Infect 23:265 e1–265 e7.

29. Fischer MA, Wamp S, Fruth A, Allerberger F, Flieger A, Halbedel S. 2020. Population structure-guided profiling of antibiotic resistance patterns in clinical *Listeria monocytogenes* isolates from Germany identifies *pbpB3* alleles associated with low levels of cephalosporin resistance. Emerg Microbes Infect 9:1804–1813.

30. Lüth S, Halbedel S, Rosner B, Wilking H, Holzer A, Roedel A, Dieckmann R, Vincze S, Prager R, Flieger A, Al Dahouk S, Kleta S. 2020. Backtracking and forward checking of human listeriosis clusters identified a multiclonal outbreak linked to *Listeria monocytogenes* in meat products of a single producer. Emerging Microbes & Infections doi:10.1080/22221751.2020.1784044:1-34.

31. Lachmann R, Halbedel S, Adler M, Becker N, Allerberger F, Holzer A, Boone I, Falkenhorst G, Kleta S, Al Dahouk S, Stark K, Luber P, Flieger A, Wilking H. 2021. Nationwide outbreak of invasive listeriosis associated with consumption of meat products in health care facilities, Germany, 2014-2019. Clin Microbiol Infect 27:1035 e1–1035 e5.

32. Ruppitsch W, Prager R, Halbedel S, Hyden P, Pietzka A, Huhulescu S, Lohr D, Schonberger K, Aichinger E, Hauri A, Stark K, Vygen S, Tietze E, Allerberger F, Wilking H. 2015. Ongoing outbreak of invasive listeriosis, Germany, 2012 to 2015. Euro Surveill 20.

33. Dominguez-Bernal G, Muller-Altrock S, Gonzalez-Zorn B, Scortti M, Herrmann P, Monzo HJ, Lacharme L, Kreft J, Vazquez-Boland JA. 2006. A spontaneous genomic deletion in *Listeria ivanovii* identifies LIPI-2, a species-specific pathogenicity island encoding sphingomyelinase and numerous internalins. Mol Microbiol 59:415–32.

34. Yin Y, Yao H, Doijad S, Kong S, Shen Y, Cai X, Tan W, Wang Y, Feng Y, Ling Z, Wang G, Hu Y, Lian K, Sun X, Liu Y, Wang C, Jiao K, Liu G, Song R, Chen X, Pan Z, Loessner MJ, Chakraborty T, Jiao X. 2019. A hybrid sub-lineage of *Listeria monocytogenes* comprising hypervirulent isolates. Nat Commun 10:4283.

35. Clayton EM, Daly KM, Guinane CM, Hill C, Cotter PD, Ross PR. 2014. Atypical *Listeria innocua* strains possess an intact LIPI-3. BMC Microbiol 14:58.

36. Ryan S, Begley M, Hill C, Gahan CG. 2010. A five-gene stress survival islet (SSI-1) that contributes to the growth of *Listeria monocytogenes* in suboptimal conditions. J Appl Microbiol 109:984–95.

37. Lee S, Ward TJ, Jima DD, Parsons C, Kathariou S. 2017. The Arsenic Resistance-Associated *Listeria* Genomic Island LGI2 Exhibits Sequence and Integration Site Diversity and a Propensity for Three *Listeria monocytogenes* Clones with Enhanced Virulence. Appl Environ Microbiol 83.

38. Parsons C, Lee S, Jayeola V, Kathariou S. 2017. Novel Cadmium Resistance Determinant in *Listeria monocytogenes*. Appl Environ Microbiol 83.

39. Müller A, Rychli K, Muhterem-Uyar M, Zaiser A, Stessl B, Guinane CM, Cotter PD, Wagner M, Schmitz-Esser S. 2013. Tn6188 - a novel transposon in *Listeria monocytogenes* responsible for tolerance to benzalkonium chloride. PLoS One 8:e76835.

40. Fischer MA, Thürmer A, Flieger A, Halbedel S. 2021. Complete Genome Sequences of Three Clinical *Listeria monocytogenes* Sequence Type 8 Strains from Recent German Listeriosis Outbreaks. Microbiol Resour Announc 10.

41. Halbedel S, Prager R, Banerji S, Kleta S, Trost E, Nishanth G, Alles G, Holzel C, Schlesiger F, Pietzka A, Schlüter D, Flieger A. 2019. A *Listeria monocytogenes* ST2 clone lacking chitinase ChiB from an outbreak of non-invasive gastroenteritis. Emerg Microbes Infect 8:17–28.

42. Halbedel S, Sperle I, Lachmann R, Kleta S, Fischer MA, Wamp S, Holzer A, Luth S, Murr L, Freitag C, Espenhain L, Stephan R, Pietzka A, Schjorring S, Bloemberg G, Wenning M, Al Dahouk S, Wilking H, Flieger A. 2023. Large Multicountry Outbreak of Invasive Listeriosis by a *Listeria monocytogenes* ST394 Clone Linked to Smoked Rainbow Trout, 2020 to 2021. Microbiol Spectr doi:10.1128/spectrum.03520-22:e0352022.

43. Ling Z, Zhao D, Xie X, Yao H, Wang Y, Kong S, Chen X, Pan Z, Jiao X, Yin Y. 2021. *inlF* Enhances *Listeria monocytogenes* Early-Stage Infection by Inhibiting the Inflammatory Response. Front Cell Infect Microbiol 11:748461.

44. Fischer MA, Engelgeh T, Rothe P, Fuchs S, Thürmer A, Halbedel S. 2022. *Listeria monocytogenes* genes supporting growth under standard laboratory cultivation conditions and during macrophage infection. Genome Res 32:1711–26.

45. Eugster MR, Haug MC, Huwiler SG, Loessner MJ. 2011. The cell wall binding domain of *Listeria* bacteriophage endolysin PlyP35 recognizes terminal GlcNAc residues in cell wall teichoic acid. Mol Microbiol 81:1419–32.

46. Rismondo J, Schulz LM, Yacoub M, Wadhawan A, Hoppert M, Dionne MS, Grundling A. 2021. EslB Is Required for Cell Wall Biosynthesis and Modification in *Listeria monocytogenes*. J Bacteriol 203.

47. Ragon M, Wirth T, Hollandt F, Lavenir R, Lecuit M, Le Monnier A, Brisse S. 2008. A new perspective on *Listeria monocytogenes* evolution. PLoS Pathog 4:e1000146.

48. Moura A, Criscuolo A, Pouseele H, Maury MM, Leclercq A, Tarr C, Bjorkman JT, Dallman T, Reimer A, Enouf V, Larsonneur E, Carleton H, Bracq-Dieye H, Katz LS, Jones L, Touchon M, Tourdjman M, Walker M, Stroika S, Cantinelli T, Chenal-Francisque V, Kucerova Z, Rocha EP, Nadon C, Grant K, Nielsen EM, Pot B, Gerner-Smidt P, Lecuit M, Brisse S. 2016. Whole genome-based population biology and epidemiological surveillance of *Listeria monocytogenes*. Nat Microbiol 2:16185.

50. Moura A, Lefrancq N, Leclercq A, Wirth T, Borges V, Gilpin B, Dallman TJ, Frey J, Franz E, Nielsen EM, Thomas J, Pightling A, Howden BP, Tarr CL, Gerner-Smidt P, Cauchemez S, Salje H, Brisse S, Lecuit M. 2020. Emergence and global spread of *Listeria monocytogenes* main clinical clonal complex. bioRxiv doi:10.1101/2020.12.18.423387:2020.12.18.423387.

50. Nuesch-Inderbinen M, Bloemberg GV, Muller A, Stevens MJA, Cernela N, Kolloffel B, Stephan R. 2021. Listeriosis Caused by Persistence of *Listeria monocytogenes* Serotype 4b Sequence Type 6 in Cheese Production Environment. Emerg Infect Dis 27:284–288.

51. McLauchlin J, Aird H, Amar C, Barker C, Dallman T, Lai S, Painset A, Willis C. 2021. An outbreak of human listeriosis associated with frozen sweet corn consumption: Investigations in the UK. Int J Food Microbiol 338:108994.

52. Vazquez-Boland JA, Wagner M, Scortti M. 2020. Why Are Some *Listeria monocytogenes* Genotypes More Likely To Cause Invasive (Brain, Placental) Infection? mBio 11.

53. Wagner E, Zaiser A, Leitner R, Quijada NM, Pracser N, Pietzka A, Ruppitsch W, Schmitz-Esser S, Wagner M, Rychli K. 2020. Virulence characterization and comparative genomics of *Listeria monocytogenes* sequence type 155 strains. BMC Genomics 21:847.

54. Wagner E, Fagerlund A, Thalguter S, Jensen MR, Heir E, Moretro T, Moen B, Langsrud S, Rychli K. 2022. Deciphering the virulence potential of *Listeria monocytogenes* in the Norwegian meat and salmon processing industry by combining whole genome sequencing and in vitro data. Int J Food Microbiol 383:109962.

55. Papic B, Pate M, Felix B, Kusar D. 2019. Genetic diversity of *Listeria monocytogenes* strains in ruminant abortion and rhombencephalitis cases in comparison with the natural environment. BMC Microbiol 19:299.

56. Cardenas-Alvarez MX, Townsend Ramsett MK, Malekmohammadi S, Bergholz TM. 2019. Evidence of hypervirulence in *Listeria monocytogenes* clonal complex 14. J Med Microbiol 68:1677–1685.

57. Ghosh P, Halvorsen EM, Ammendolia DA, Mor-Vaknin N, O’Riordan MXD, Brumell JH, Markovitz DM, Higgins DE. 2018. Invasion of the Brain by *Listeria monocytogenes* Is Mediated by InlF and Host Cell Vimentin. mBio 9.

58. Faralla C, Rizzuto GA, Lowe DE, Kim B, Cooke C, Shiow LR, Bakardjiev AI. 2016. InlP, a New Virulence Factor with Strong Placental Tropism. Infect Immun 84:3584–3596.

59. Balogh D, Eckel K, Fetzer C, Sieber SA. 2022. *Listeria monocytogenes* utilizes the ClpP1/2 proteolytic machinery for fine-tuned substrate degradation at elevated temperatures. RSC Chem Biol 3:955–971.

60. Elsholz AKW, Birk MS, Charpentier E, Turgay K. 2017. Functional Diversity of AAA+ Protease Complexes in *Bacillus subtilis*. Front Mol Biosci 4:44.

61. Elfmann C, Zhu B, Stülke J, Halbedel S. 2023. ListiWiki: A database for the foodborne pathogen *Listeria monocytogenes*. Int J Med Microbiol 313:151591.

62. Glaser P, Frangeul L, Buchrieser C, Rusniok C, Amend A, Baquero F, Berche P, Bloecker H, Brandt P, Chakraborty T, Charbit A, Chetouani F, Couve E, de Daruvar A, Dehoux P, Domann E, Dominguez-Bernal G, Duchaud E, Durant L, Dussurget O, Entian KD, Fsihi H, Garcia-del Portillo F, Garrido P, Gautier L, Goebel W, Gomez-Lopez N, Hain T, Hauf J, Jackson D, Jones LM, Kaerst U, Kreft J, Kuhn M, Kunst F, Kurapkat G, Madueno E, Maitournam A, Vicente JM, Ng E, Nedjari H, Nordsiek G, Novella S, de Pablos B, Perez-Diaz JC, Purcell R, Remmel B, Rose M, Schlueter T, Simoes N, et al. 2001. Comparative genomics of *Listeria* species. Science 294:849–52.

63. Mandin P, Repoila F, Vergassola M, Geissmann T, Cossart P. 2007. Identification of new noncoding RNAs in *Listeria monocytogenes* and prediction of mRNA targets. Nucleic Acids Res 35:962–74.

64. Halbedel S, Hahn B, Daniel RA, Flieger A. 2012. DivIVA affects secretion of virulence-related autolysins in *Listeria monocytogenes*. Mol Microbiol 83:821–39.

65. Rismondo J, Bender JK, Halbedel S. 2017. Suppressor Mutations Linking *gpsB* with the First Committed Step of Peptidoglycan Biosynthesis in *Listeria monocytogenes*. J Bacteriol 199.

66. Köser CU, Fraser LJ, Ioannou A, Becq J, Ellington MJ, Holden MT, Reuter S, Torok ME, Bentley SD, Parkhill J, Gormley NA, Smith GP, Peacock SJ. 2014. Rapid single-colony whole-genome sequencing of bacterial pathogens. J Antimicrob Chemother 69:1275–81.

67. Ruppitsch W, Pietzka A, Prior K, Bletz S, Fernandez HL, Allerberger F, Harmsen D, Mellmann A. 2015. Defining and Evaluating a Core Genome Multilocus Sequence Typing Scheme for Whole-Genome Sequence-Based Typing of *Listeria monocytogenes*. J Clin Microbiol 53:2869–76.

68. Letunic I, Bork P. 2021. Interactive Tree Of Life (iTOL) v5: an online tool for phylogenetic tree display and annotation. Nucleic Acids Res 49:W293–W296.

69. Deeks JJ, Higgins JP, Altman DG, Group obotCSM. 2019. Analysing data and undertaking meta-analyses, p 241-284, Cochrane Handbook for Systematic Reviews of Interventions 10.1002/9781119536604.ch10.

71. Wick R, Menzel P. 2021. Filtlong. https://github.com/rrwick/Filtlong. Accessed 01.07.2021.

71. Wick RR, Judd LM, Gorrie CL, Holt KE. 2017. Unicycler: Resolving bacterial genome assemblies from short and long sequencing reads. PLoS Comput Biol 13:e1005595.

72. Becker L, Fuchs S, Pfeifer Y, Semmler T, Eckmanns T, Korr G, Sissolak D, Friedrichs M, Zill E, Tung M-L, Dohle C, Kaase M, Gatermann S, Rüssmann H, Steglich M, Haller S, Werner G. 2018. Whole Genome Sequence Analysis of CTX-M-15 Producing *Klebsiella* Isolates Allowed Dissecting a Polyclonal Outbreak Scenario. Frontiers in Microbiology 9.

73. Abdelhamed H, Lawrence ML, Karsi A. 2015. A novel suicide plasmid for efficient gene mutation in *Listeria monocytogenes*. Plasmid 81:1–8.

