## Supplementary Tables S2-S3 for "High density genomic surveillance and risk profiling of clinical *Listeria monocytogenes* subtypes in Germany, 2018-2021"

**Tab. S2:** Genomic features of representative isolates from selected German outbreak clusters.

| | **Alpha10** | **Epsilon1b** | **Theta3a** | **Kappa8** | **Eta7** |
|---|---|---|---|---|---|
| strain | 21-03201 | 11-04869 | 16-02236 | 19-07394 | 19-02390 |
| source type | clinical isolate | clinical isolate | clinical isolate | clinical isolate | clinical isolate |
| isolation source | not reported | blood | CSF[1] | blood | nasal swap |
| year of isolation | 2021 | 2011 | 2016 | 2019 | 2019 |
| NCBI accession | CP111149 | CP110922 | CP111148 | CP113891 | CP111150 |
| ENA accession | ERS14238016 | ERS2103006 | ERS14238017 | ERS14291506 | ERS14238018 |
| phyl. lineage | I | I | I | I | I |
| PCR serogroup | IVb | IVb | IVb | IVb | IVb |
| ST/CC[1] | ST1/CC1 | ST6/CC6 | ST249/CC315 | ST1/CC1 | ST6/CC6 |
| CT[2] Ruppitsch | CT6329 | CT90 | CT4449 | CT4961 | CT7504 |
| CT[2] Pasteur | CT9528 | CT443 | CT12857 | CT3989 | CT7198 |
| sequencing method | NextSeq/MinION | MiSeq/MinION | MiSeq/MinION | MiSeq/MinION | MiSeq/MinION |
| long read raw data | ERR10513163 | ERR10513160 | ERR10513161 | ERR10556194 | ERR10513162 |
| short read raw data | ERR10481070 | ERR2261336 | ERR10481069 | ERR10556193 | ERR10481068 |
| genome size | 2,946,483 bp | 3,034,959 bp | 2,976,730 bp | 2,947,414 bp | 2,956,565 bp |
| G/C content | 38.0% | 38.0% | 38.0% | 38.0% | 38.0% |
| CDS[3] (protein) | 2,850 | 2,988 | 2,851 | 2,852 | 2,860 |
| rRNA operons | 6 | 6 | 6 | 6 | 6 |
| tRNA genes | 67 | 67 | 67 | 67 | 65 |
| PAIs[3] | LIPI-1, LIPI-3 | LIPI-1, LIPI-3 | LIPI-1, LIPI-4 | LIPI-1, LIPI-3 | LIPI-1, LIPI-3 |
| SSIs[3] | - | - | SSI-1 | - | - |
| plasmids | - | pLMST6 (CP110923) | - | - | - |
| intact prophages[4] | 1 | 3 | 1 | 1 | 1 |
| insertion sites | *comK* | tRNA$^{Lys}$ (*lmot01*)<br>OQ358_06820<br>*rpsI* | tRNA$^{Arg}$ (*lmot17*) | tRNA$^{Lys}$ (*lmot01*) | *tsf* |
| PMSCs[3] | ORY88_00790 (*lmo0140*)<br>ORY88_10650 (*lmo2084*) | OQ358_01125 (*lmo0140*)<br>OQ358_02035 (*lmo0310*)<br>OQ358_02370 (*lmo0380*)<br>OQ358_02740 (*lmo0440*) | ORY89_02055 (*lmo0380*)<br>ORY89_07620 (*ispG*)<br>ORY89_12510 (*lmo2409*) | OZX46_01155 (*lmo0140*)<br>OZX46_03895 (*lmo0671*)<br>OZX46_11015 (*lmo2084*)<br>OZX46_14505 (*lmo2797*) | ORY90_00355 (*essC*)<br>ORY90_00795 (*lmo0140*)<br>ORY90_01705 (*lmo0310*)<br>ORY90_02040 (*lmo0380*)<br>ORY90_02410 (*lmo0440*)<br>ORY90_03140 (*lmo0595*)<br>ORY90_05765 (*lmo1123*)<br>ORY90_05820 (*lmo1134*)<br>ORY90_14515 (*lmo2781*) |

[1] ST - sequence type (ST) and clonal complex (CC) according to seven locus MLST (1),
[2] CT - complex types according to Ruppitsch and Pasteur cgMLST schemes (2, 3),
[3] abbreviations: CSF - cerebrospinal fluid, CDS – coding sequences, PAI – pathogenicity associated island, PMSC – premature stop codon, SSI – stress survival islet,
[4] according to PHASTER (4).

**Tab. S3:** Subpopulations of clinical *L. monocytogenes* isolates with inactivating mutations in selected genes.

| gene | name | function | mutation | affected subgroup[1] | expected phenotype | number of isolates[2] |
|---|---|---|---|---|---|---|
| *lmo0105* | *chiB* | chitinase | S345X | Iota6 (CT6520, ST4, IVb) | no chitin utilization (5) | 2/2 |
| | | | | Iota7 (CT8189, ST4, IVb) | virulence defect in mice (6) | 2/2 |
| | | | | Tau4 (CT6460, ST4, IVb) | | 5/5 |
| *lmo0409* | *inlF* | internalin F | Q451X | Tau8 (CT9031, ST451, IIa) | host cell entry defect (7) | 16/16 |
| *lmo0433* | *inlA* | internalin A | E326X | My5 (CT6466, ST9, IIc) | host cell entry defect (8) | 2/2 |
| | | | Q492X | ST121 (IIa) | | 15/15 |
| | | | N546fs | Gamma11 (CT1698, ST9, IIc) | | 2/2 |
| | | | | Rho3 (CT1690, ST9, IIc) | | 3/4 |
| *lmo1076* | *aut* | Auto autolysin | K132X | Psi2a/b (CT1234, ST155, IIa) | host cell entry defect (9) | 3/3 |
| | | | S144X | Omikron11 (CT14992, ST155, IIa) | | 2/2 |
| | | | W163X | Delta8 (CT4295, ST121, IIa) | | 4/4 |
| | | | N189fs | Pi5 (CT6665, ST3, IIb) | | 5/5 |
| | | | N339fs | Alpha5 (CT6583, ST3, IIb) | | 3/3 |
| *lmo1138* | *clpP1* | accessory ClpP1 protease | L34X | ST16 (IIa) | protein turnover defect (10) | 9/9 |
| *lmo1412* | *flaR* | required for flagellin expression | W50X | ST3 (IIb) | motility defect (11) | 25/29 |
| | | | D67fs | ST29 (IIa) | | 9/11 |
| | | | | ST37 (IIa) | | 58/64 |
| | | | | ST427 (IIa) | | 7/9 |
| | | | | ST1344 (IIa) | | 2/2 |
| | | | K75fs | ST18 (IIa) | | 13/13 |
| | | | | ST26 (IIa) | | 13/15 |
| | | | | ST200 (IIb) | | 5/5 |
| | | | | ST207 (IIa) | | 3/4 |
| *lmo1441* | *ispG* | hydroxymethylbutenyl diphosphate synthase | R115X | ST249 (IVb) | virulence defect in mice (12) | 24/24 |
| *lmo2467* | | chitin binding protein | S82X | My5 (CT6466, ST9, IIc) | virulence defect in mice (6) | 2/2 |
| | | | W224X | ST29 (IIa) | | 8/11 |
| *lmo2550* | *csbB* | required for WTA decoration with GlcNAc | A81fs | Sigma5 (CT5715, ST504, IIa) | GlcNAc-less WTA (13) | 12/12 |
| | | | L104fs | Omega5 (CT1138, ST87, IIb) | | 3/11 |
| *lmo2769* | *eslA* | ABC transporter, ATP-binding protein | I160fs | ST38 (IIa) | lysozyme sensitive (14) | 3/3 |
| | | | I229fs | ST427 (IIa) | | 9/9 |

[1] STs associated with reduced MFL and NL risks were marked by a double underscore, those associated with reduced NL or MFL risk with a single underscore.

[2] Only mutations, which were found at least twice per phylogroup, were included.

### References


1. Ragon M, Wirth T, Hollandt F, Lavenir R, Lecuit M, Le Monnier A, Brisse S. 2008. A new perspective on *Listeria monocytogenes* evolution. PLoS Pathog 4:e1000146.
2. Ruppitsch W, Pietzka A, Prior K, Bletz S, Fernandez HL, Allerberger F, Harmsen D, Mellmann A. 2015. Defining and Evaluating a Core Genome Multilocus Sequence Typing Scheme for Whole-Genome Sequence-Based Typing of *Listeria monocytogenes*. J Clin Microbiol 53:2869-76.
3. Moura A, Tourdjman M, Leclercq A, Hamelin E, Laurent E, Fredriksen N, Van Cauteren D, Bracq-Dieye H, Thouvenot P, Vales G, Tessaud-Rita N, Maury MM, Alexandru A, Criscuolo A, Quevillon E, Donguy MP, Enouf V, de Valk H, Brisse S, Lecuit M. 2017. Real-Time Whole-Genome Sequencing for Surveillance of *Listeria monocytogenes*, France. Emerg Infect Dis 23:1462-1470.
4. Arndt D, Grant JR, Marcu A, Sajed T, Pon A, Liang Y, Wishart DS. 2016. PHASTER: a better, faster version of the PHAST phage search tool. Nucleic Acids Res 44:W16-21.
5. Leisner JJ, Larsen MH, Jorgensen RL, Brondsted L, Thomsen LE, Ingmer H. 2008. Chitin hydrolysis by *Listeria* spp., including *L. monocytogenes*. Appl Environ Microbiol 74:3823-30.
6. Chaudhuri S, Bruno JC, Alonzo F, 3rd, Xayarath B, Cianciotto NP, Freitag NE. 2010. Contribution of chitinases to *Listeria monocytogenes* pathogenesis. Appl Environ Microbiol 76:7302-5.
7. Ling Z, Zhao D, Xie X, Yao H, Wang Y, Kong S, Chen X, Pan Z, Jiao X, Yin Y. 2021. *inlF* Enhances *Listeria monocytogenes* Early-Stage Infection by Inhibiting the Inflammatory Response. Front Cell Infect Microbiol 11:748461.
8. Gaillard JL, Berche P, Frehel C, Gouin E, Cossart P. 1991. Entry of *L. monocytogenes* into cells is mediated by internalin, a repeat protein reminiscent of surface antigens from gram-positive cocci. Cell 65:1127-41.
9. Cabanes D, Dussurget O, Dehoux P, Cossart P. 2004. Auto, a surface associated autolysin of *Listeria monocytogenes* required for entry into eukaryotic cells and virulence. Mol Microbiol 51:1601-14.
10. Balogh D, Eckel K, Fetzer C, Sieber SA. 2022. *Listeria monocytogenes* utilizes the ClpP1/2 proteolytic machinery for fine-tuned substrate degradation at elevated temperatures. RSC Chem Biol 3:955-971.
11. Sanchez-Campillo M, Dramsi S, Gomez-Gomez JM, Michel E, Dehoux P, Cossart P, Baquero F, Perez-Diaz JC. 1995. Modulation of DNA topology by *flaR*, a new gene from *Listeria monocytogenes*. Mol Microbiol 18:801-11.
12. Heuston S, Begley M, Davey MS, Eberl M, Casey PG, Hill C, Gahan CGM. 2012. HmgR, a key enzyme in the mevalonate pathway for isoprenoid biosynthesis, is essential for growth of Listeria monocytogenes EGDe. Microbiology (Reading) 158:1684-1693.
13. Eugster MR, Haug MC, Huwiler SG, Loessner MJ. 2011. The cell wall binding domain of *Listeria* bacteriophage endolysin PlyP35 recognizes terminal GlcNAc residues in cell wall teichoic acid. Mol Microbiol 81:1419-32.
14. Rismondo J, Schulz LM, Yacoub M, Wadhawan A, Hoppert M, Dionne MS, Grundling A. 2021. EslB Is Required for Cell Wall Biosynthesis and Modification in *Listeria monocytogenes*. J Bacteriol 203.
