## Supplementary Figures S1-S4 for "High density genomic surveillance and risk profiling of clinical *Listeria monocytogenes* subtypes in Germany, 2018-2021"

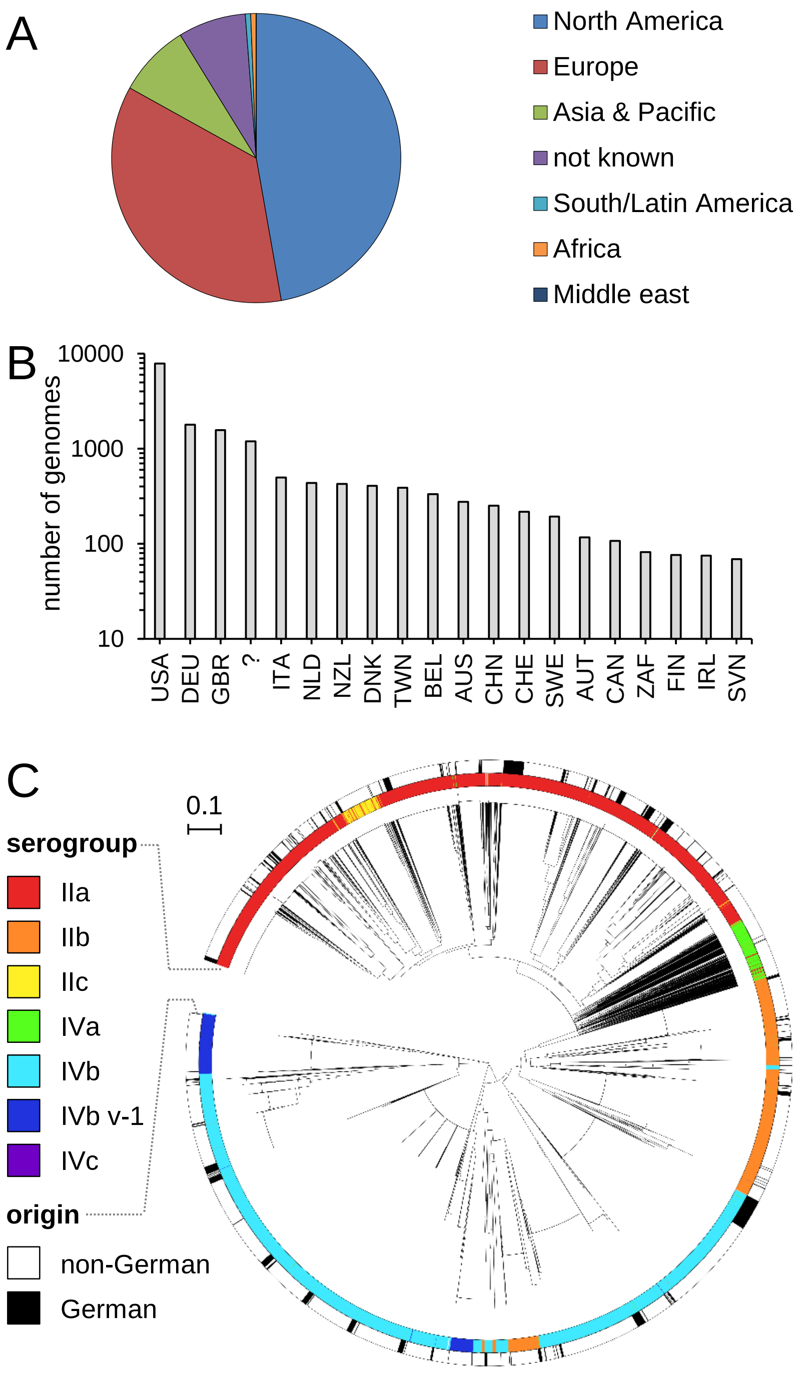


**Fig. S1: Comparison of the population structures of clinical *L. monocytogenes* isolates from in- and outside Germany.**

(A) Representation of the different world regions in the dataset used for comparison. 15,155 non-German NCBI isolate genomes and 1,802 isolate genomes from Germany are included.

(B) Diagram illustrating the countries of isolation in the same dataset. Countries with the most isolates are shown, ? – country of isolation not known.

(C) Neighbor joining tree calculated on seven locus MLST data showing the phylogenetic structure of the 15,155 non-German and the 1,802 German clinical isolates. Tree tips are colored according to molecular PCR serogroups (inner ring) and to their geographic origin (outer ring).

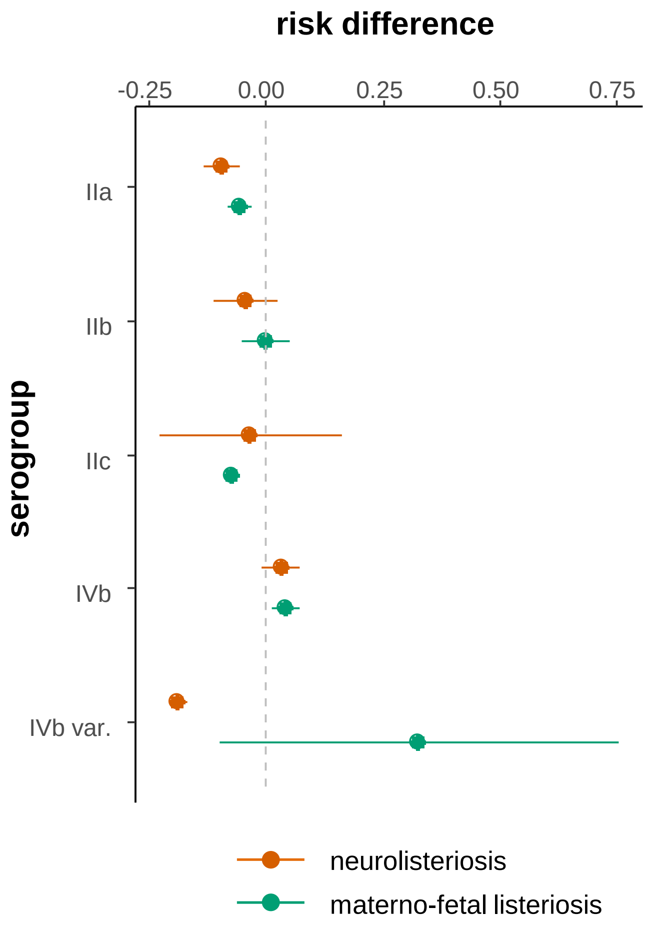


**Fig. S2: Clinical disease manifestation as reported during case notification.**
Risks for NL and MFL manifestations are expressed as risk differences together with 95% confidence intervals for the individual serogroups. Data were calculated based on 1,323 isolate/notification case pairs, for which information on the disease manifestation were available.

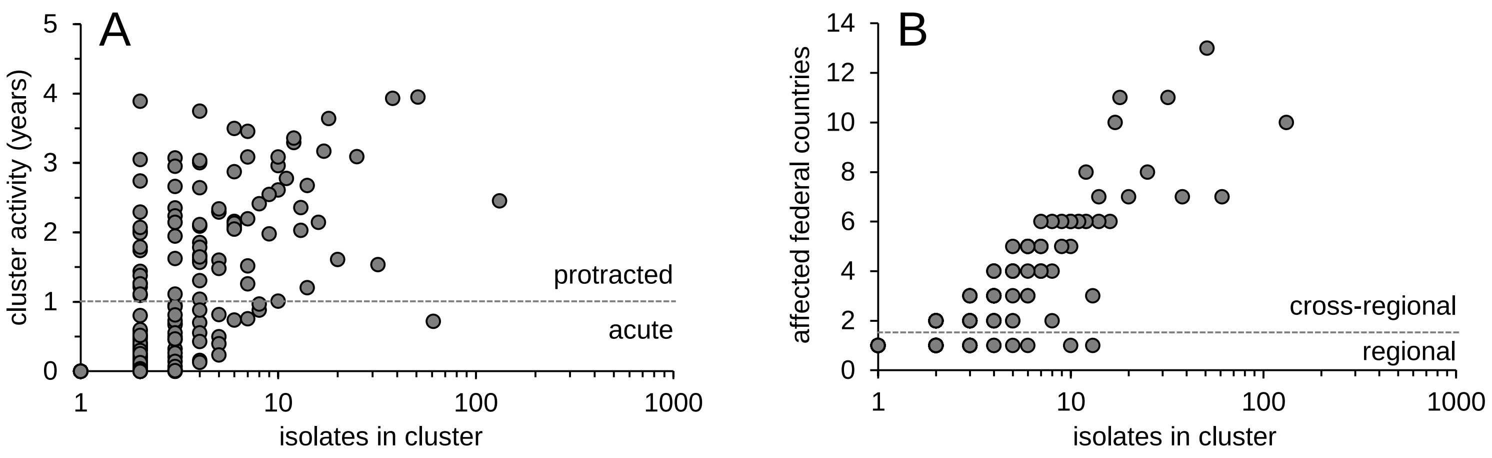


**Fig. S3: Duration and geographical spread of listeriosis clusters.**
(A) Diagram showing the duration of all listeriosis clusters within the four-year study period plotted against the number of isolates.
(B) Diagram illustrating the geographical distribution by plotting the number of involved federal countries in Germany against cluster size.

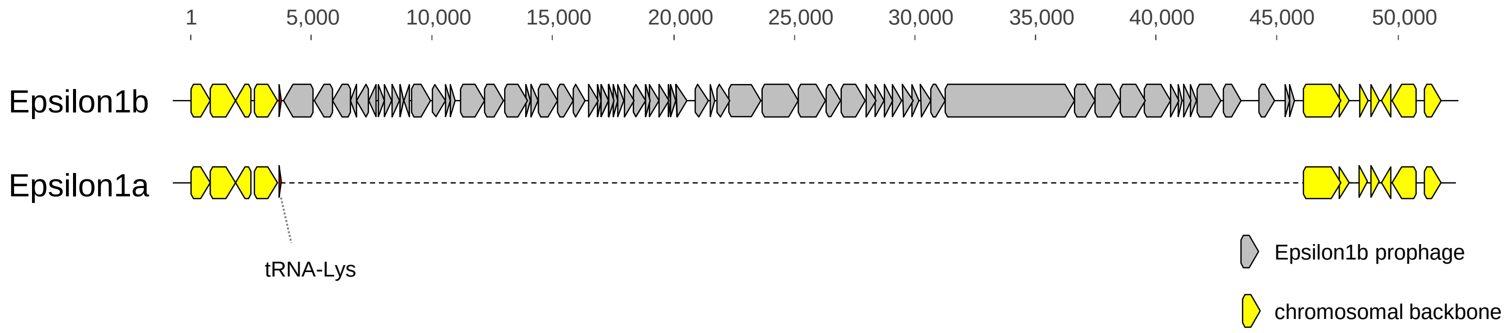


**Fig. S4:** Absence of an Epsilon1b prophage in the *L. monocytogenes* Epsilon1a outbreak clone
Comparison of the tRNA-Lys chromosomal regions of strains 18-04540 (Epsilon1a, accession number: CP063383) and 11-04869 (Epsilon1b, accession number: CP110922).
